# Haplotype-resolved assemblies and variant benchmark of a Chinese Quartet

**DOI:** 10.1101/2022.09.08.504083

**Authors:** Peng Jia, Lianhua Dong, Xiaofei Yang, Bo Wang, Tingjie Wang, Jiadong Lin, Songbo Wang, Xixi Zhao, Tun Xu, Yizhuo Che, Ningxin Dang, Luyao Ren, Yujing Zhang, Xia Wang, Fan Liang, Yang Wang, Jue Ruan, The Quartet Project Team, Yuanting Zheng, Leming Shi, Jing Wang, Kai Ye

**Author notes:** These authors contributed equally (Peng Jia, Lianhua Dong). To whom correspondence should be addressed. (Ye K), (Wang J).

## Abstract

As the state-of-the-art sequencing technologies and computational methods enable investigation of challenging regions in the human genome, an update variant benchmark is demanded. Herein, we sequenced a Chinese Quartet, consisting of two monozygotic twin daughters and their biological parents, with multiple advanced sequencing platforms, including Illumina, BGI, PacBio, and Oxford Nanopore Technology. We phased the long reads of the monozygotic twin daughters into paternal and maternal haplotypes using the parent-child genetic map. For each haplotype, we utilized advanced long reads to generate haplotype-resolved assemblies (HRAs) with high accuracy, completeness, and continuity. Based on the ingenious quartet samples, novel computational methods, high-quality sequencing reads, and HRAs, we established a comprehensive variant benchmark, including 3,883,283 SNVs, 859,256 Indels, 9,678 large deletions, 15,324 large insertions, 40 inversions, and 31 complex structural variants shared between the monozygotic twin daughters. In particular, the preciously excluded regions, such as repeat regions and the human leukocyte antigen (HLA) region, were systematically examined. Finally, we illustrated how the sequencing depth correlated with the *de novo* assembly and variant detection, from which we learned that 30 × HiFi is a balance between performance and cost. In summary, this study provides high-quality haplotype-resolved assemblies and a variant benchmark for two Chinese monozygotic twin samples. The benchmark expanded the regions of the previous report and adapted to the evolving sequencing technologies and computational methods.

## Background

Since the dawn of the genome era, genomic variations, including single nucleotide variations (SNVs), small insertions/deletions (Indels) and structural variants (SVs), have been extensively detected and proved to contribute to many diseases, such as Mendelian disorders and cancers^1–4^. Thus, authoritative and comprehensive variant benchmarks are crucial for precisely understanding genetic variations in clinical samples. Many variant benchmarks and genomic reference materials have been established for the community to evaluate their variant detection pipelines during the past decades^5–15^. For example, the Genome in a Bottle (GIAB) Consortium developed seven reference materials and high-confidence benchmarks for both small variants^10^ and structural variants^7^, prompting the pipeline evaluation in genomic analysis. Another companion study released a robust benchmark on the Certified Reference Materials for whole genome-variant assessment to reveal the variant detection biases among different short-read sequencing platforms and among sequencing centers^15^. Nevertheless, these studies focus on the simple variant types and high-confidence regions for short reads, ignoring the complex regions and complex variant types that are accessible for long read sequencing technologies.

Advanced sequencing technologies^16–18^, including PacBio HiFi and Oxford Nanopore ultra-long reads, were recently leveraged to assemble a complete hydatidiform mole (CHM13) at telomere-to-telomere levels^19^, making it possible to resolve many medical-related genes and regions excluded by previous benchmarks. Another remarkable investigation of genetic variants by the Human Genome Structural Variation Consortium (HGSVC) demonstrates that high-quality haplotype-resolved assemblies (HARs) detect more variants than previous read-alignment-based strategies^20^. Based on the high-quality HRAs, variants located in complex regions, such as simple repeat (SR), segmental duplication (SD), variable number tandem repeat (VNTR), and short tandem repeat (STR), were resolved. In addition to high-quality reads and assemblies, novel computational methods such as Sniffles^21^, cuteSV^22^, and SVision^23^ were also developed to reveal complex SVs in the human genome.

As samples for benchmarking in practices, a single sample or even a trio is difficult to deal with the random variants induced by contamination in cell line culture and transportation^24^. To address this problem, we included a “Chinese Quartet”, consisting of two monozygotic twin daughters (LCL5 and LCL6) and their biological parents (LCL7 and LCL8), in this study. Notably, the DNA of four samples was approved as Certified Reference Materials (CRMs) for whole genome-variant assessment (GBW09900~GBW09903) by the State Administration for Market Regulation in China. We applied advanced sequencing technologies to the four samples and emphatically assembled high-quality haplotype-resolved genomes for the monozygotic twins. We demonstrated that two haplotypes of the diploid samples achieved high performance in terms of accuracy, continuity, and completeness. Benefitting from the ingenious samples, the advanced sequencing technologies, high-quality HRAs, and novel computational methods, we construct a comprehensive benchmark for all scales of variants. In particular, we extend the variant benchmark to complex regions and complex variant types.

## Results

### Sample processing and sequencing

In this study, we included various sequencing data of the Chinese Quartet, parents and monozygotic twin daughters, to construct a high-quality genome and variant benchmark for the Chinese Han population. To obtain high-quality assemblies for the twin daughters, we generated ~ 50 × HiFi (read length N50 = 13~14 kb), ~ 100 × ONT regular (read length N50 = 20~25 kb) reads for each of four samples, and addition ~ 30 × ONT ultra-long (read length N50 = 77 kb) reads for one twin sample, LCL5 (Table S1). To establish a robust variant benchmark for the twin daughters, we used ~ 160 × Illumina (150bp read length) and ~100 × BGI (100bp read length) reads and a variety of long reads to discover and evaluate the variants shared between the monozygotic twin daughters (Table S1).

### Haplotype-resolved genome assembly

Since monozygotic twins are generally considered genetically identical with limited somatic substations^25^, we first merged the data of these two samples and endeavored to generate high-quality haplotype-resolved genomes. We phased HiFi, ONT regular, and ONT ultra-long reads of the monozygotic twins into paternal (CQ-P) and maternal (CQ-M) haplotypes and assembled each haplotype using a hybrid assembly strategy (Fig. S1). First, high-quality SNVs and Indels were obtained from a previous study^13^, and both the sharing patterns among trios and their concurrence on HiFi reads^26^. Next, long reads including HiFi, ONT regular, and ONT ultra-long of two twin daughters were separated into two haplotypes with the phased variants^26^. Overall, we phased 76.2 % of HiFi reads, 65.0 % of ONT regular reads, and 72.8 % of ONT ultra-long reads, and the unphased reads were assigned to the two haplotypes randomly (Table S2). For each haplotype of the two twin daughters, we obtained around 53 × HiFi, 95 × ONT regular, and 14 × ONT ultra-long reads (Table S2). We assembled ONT reads using shasta^27^ and flye^28^ and assembled HiFi reads using hifiasm^29^, hicanu^30^, and flye^28^, yielding five haplotype-resolved assemblies (Table S3). After that, the hifiasm contigs were scaffolded using ragtag^31^ and the other four assemblies were used to fill the gaps in the hifiasm scaffolds (see methods and Supplementary Notes). Finally, the two haplotypes of twin daughters were further polished with phased HiFi reads^32^ (see methods and Supplementary Notes).

The final two haplotypes contained 297 contigs and 276 for CQ-P and CQ-M, respectively, and both haplotypes had a length of around 3.05 Gb. The contig N50 values of two haplotypes are ~ 132M, about 2-fold of GRCh38.p13, suggesting high continuousness of the obtained phased assemblies compared to previous reports^33–37^ (Table 1, S3 and S4). Notably, seven and nine chromosomes of two haplotypes were gap free. Meanwhile, 20 and 18 chromosome arms in CQ-P and CQ-M were successfully represented as a single contig, respectively (Fig. S2, S3, and Table S5). Furthermore, CQ-P and CQ-M closed 236 and 251 gaps in GRCh38, respectively (**Fig. 1A**, and Fig. S4). For example, the HiFi read depth illustrated that GRCh38 gaps near the centromere of chromosome 17 were filled by both CQ-P and CQ-M haplotypes **(Fig. 1B)**. Another further example, a previous reported polymorphic inversion by CHM13^38^ at chromosome 8p23.1, was also identified, and the flanking gaps of the ~ 4M inversion were accurately resolved by both haplotypes (**Fig. 1C** and Fig. S5).

**Figure 1.**
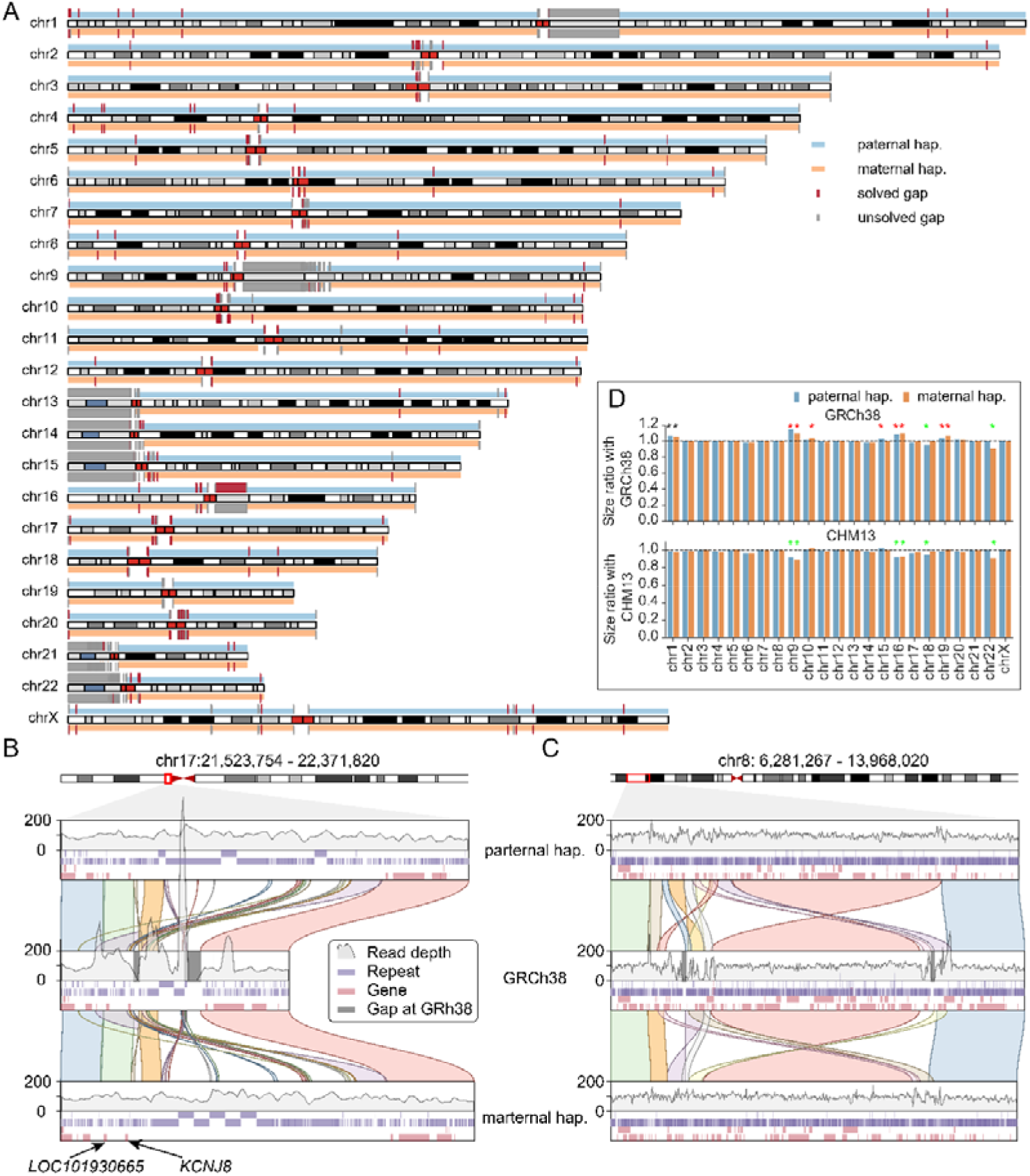
An overview of Chinese Quartet assemblies. **A** Idiogram depicts the alignments between the GRCh38 (gray rectangles) and two Chinese Quartet haplotypes (blue rectangles for CQ-P and orange for CQ-M). The red rectangles represent the GRCh38 gaps filled by Chinese Quartet assemblies, while the gray rectangles refer to unresolved gaps. **B** and **C** Examples of gaps resolved by Chinese Quartet assemblies. The top and bottom channels represent the paternal and maternal haplotypes, respectively. The middle channel represents the GRCh38. The depths of HiFi reads on three genomes are shown with gray lines. The repeat regions and genes are labeled with purple and pink rectangles, and the gaps in GRCh38 are labeled with gray rectangles. **D** The bar plots show the percentage size of Chinese Quartet assembled chromosomes relative to CHM13 (top) and GRCh38 chromosomes (bottom), without including Ns. The chromosome with more than 3% difference in length is labeled with star.

**Table 1:** Summary statistics comparison of haplotype-resolved assemblies of Chinese Quartet and other assemblies.

| Sample | Haplotype | Genome length (Gb) | No. of contigs | Contig N50 (Mb) | Completeness (BUSCO) | QV | Switch error |
| --- | --- | --- | --- | --- | --- | --- | --- |
| Chinese Quartet | Paternal | 3.05 | 279 | 132.84 | 95.7% | 50 - 58 | 0.050% |
|  | Maternal | 3.05 | 276 | 132.84 | 95.7% | 52 - 59 | 0.048% |
| HJ | Paternal | 3.07 | 1330 | 28.15 | 94.9% | 52 - 59 | 0.815% |
|  | Maternal | 2.91 | 896 | 25.90 | 93.5% | 54 - 58 | 0.813% |
| NA12878 | Hap1 | 2.88 | 4,363 | 18.3 | 95.5% | 51 - 60 | 0.449% |
|  | Hap2 | 2.88 | 4,449 | 21.9 | 95.4% | 51 - 60 | 0.435% |
| HG00733 | Hap1 | 2.92 | 3,728 | 23.7 | 94.9% | 50 - 59 | 0.169% |
|  | Hap2 | 2.92 | 3,795 | 25.9 | 95.1% | 51 - 59 | 0.171% |
| YH2.0 | Collapsed | 2.91 | 361,157 | 0.02 | 94.2% | NA | NA |
| HX1 | Collapsed | 2.93 | 5,845 | 8.33 | 94.0% | NA | NA |
| NH1.0 | Collapsed | 2.89 | 11,019 | 3.6 | 94.6% | NA | NA |
| GRCh38.p13* | Collapsed | 3.21 | 685 | 56.41 | 94.7% | NA | NA |

*Note:* \* GRCh38 without the alternative sequences; NA: not available.

We demonstrated that ten chromosomes (5 paternal and 5 maternal) of our assemblies had more than a 3% increase in length compared with GRCh38, while six chromosomes (3 paternal and 3 maternal) had a 3%decrease in length compared to CHM13 (**Fig. 1D**). To further assess the completeness of CQ-P and CQ-M, we aligned two haplotypes against GRCh38 and observed that CQ-P and CQ-M covered 97.59% and 97.55% of the GRCh38 genome, respectively (Table S6). The completeness evaluation by BUSCO^39^ (v5.1.3) showed that our phased genomes resolved 95.7% of complete genes from the mammalia_odb10 library, indicating that our assemblies were highly complete as well (Table 1).

To characterize the reference material comprehensively, we annotated genes and novel sequences of two haplotypes (Fig. S6). We found 8.4 M and 8.8 M novel sequences in CQ-M and CQ-P, respectively, when compared to GRCh38. Most novel sequences were located in centromeric and acrocentric regions (Fig. S7). To annotate our genomes, we converted the gene coordinates of GRCh38.p13 to CQ-P and CQ-M with liftoff ^40^, of which 96.62% (19207/19878) and 96.54% (19191/19878) of protein-coding genes were successfully converted. To annotate genes at novel sequences, we then masked the repeat sequences and annotated the protein-coding genes by Augustus^41^. We finally obtained 45 and 58 novel genes in CQ-P and CQ-M, respectively (Table S7). The most abundant functional domains in these novel genes included domains such as ElonginA binding-protein 1 (PF15870), Poly-adenylate binding protein domain (PF00658), Kinase suppressor of RAS, SAM-like domain (PF13543) and Extensin domain (PF04554).

### Variant benchmark construction

Since each sequencing technology and variant pipeline had its own advantages, we involved short reads, long reads, and haplotype-resolved assemblies to discover all scales of variants for the monozygotic twins (see Methods). In particular, the twin daughters were regarded as two biological replicates, so that only variants supported by both samples were kept in the final benchmark (Fig. S8 and S9, see Methods).

#### SNV and Indel benchmark construction

For SNVs and Indels, Illumina calls were downloaded from the previous study^13^, HiFi calls were generated by the minimap2-deepvariant pipeline^42, 43^. Both the Illumina and HiFi calls were filtered by read depth, allele frequency, and Mendelian rule. Meanwhile, three haplotype-resolved assemblies by HiFi reads were used for variant discovery by PAV^20^, and only variants supported by all three assemblies were included in the HRA callset (Fig. S8, see Methods).

We released 3,883,283 SNVs and 859,256 Indels for the monozygotic twins (**Fig. 2A**), of which 91.1% of SNVs and 91.8% of Indels were also observed by BGI reads (**Fig. 2B**, and Fig. 10). Notably, long-read assembly (HRAs) based variant calling strategies contributed to 97.9% (3,803,062) of SNVs and 98.4% (845,085) of Indels, while long-read HiFi mapping based approaches accounted for 93.2% (3,619,614) of SNVs and 70.1% (602,343) of Indels. Illumina short-read mapping based variant calling result yielded 81.0% (3,144,055) of SNVs and 45.1% (387,741) of Indels.

**Figure 2.**
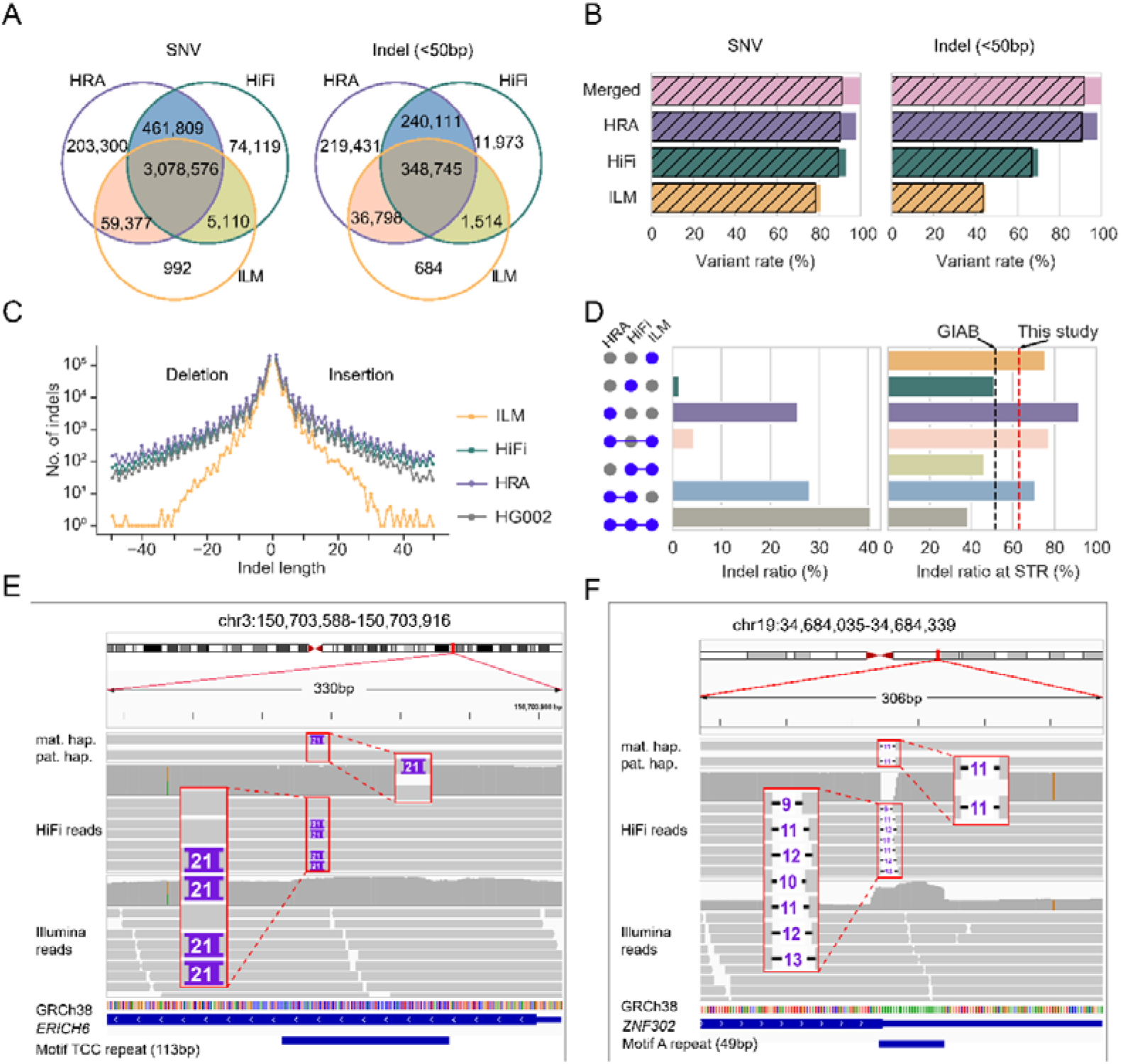
Small variant benchmark of Chinese Quartet. **A** Overlap of SNVs and Indels among ILM, HiFi, and HRA, respectively. **B** Bar plot depicts the percentage of ILM, HiFi, and HRA calls in SNV (left) and Indel (right) benchmark, with gray stripes representing the percentages of calls supported by BGI reads. **C** Indel length distribution of Indels across HG002 and three callsets of Chinese Quartet. **D** Left bar represents the percentages of indels in different combinations of three technologies. Right bar represents the ratio of Indels at STR regions across different combinations of three technologies. **E** IGV snapshot shows a heterozygous deletion at a TCC repeat. This deletion is detected by both HRA and HiFi reads. **F** IGV snapshot shows a homozygous insertion at a homopolymer region. This deletion is only detected by HRA.

As expected, the Indel length distribution demonstrated that the sensitivities of Illumina, HiFi, and HRAs to detect Indel increased accordingly (**Fig. 2C** and Fig. S11). Meanwhile, we found that HiFi and HRA detected more Indels in complex regions like STR (**Fig. 2D**, Fig. S12 and S13). In particular, HRA detected 25.5% of Indels specifically, of which 91.7% were in STR regions. For example, a 21 bp heterozygous insertion of TCC repeat at *ERICH6* was accurately identified by both HRAs and HiFi reads, but missed by Illumina data due to its shorter read length (**Fig. 2E**). Another example was that an 11bp deletion in a homopolymer region (49 bp A repeat) of *ZNF302* was missed by both HiFi and Illumina reads but detected by HRAs, indicating the vantage of HRAs for Indel detection in homopolymer regions (**Fig. 2F**).

#### Large deletion and insertion benchmark construction

Structural variants affected more nucleotides and were more deleterious than SNVs and Indels^3^, although they are relatively rare compared to SNVs and Indels. However, SV detection and benchmarking remain challenging. To overcome the biases of SV detection across different technologies, SVs from Illumina reads, HiFi reads, and haplotype-resolved assemblies were discovered, filtered, and merged. Illumina calls were generated by four prevalent callers, including Manta^44^ (v1.6.0), Delly^45^ (v0.9.1), Lumpy^46^ (v0.2.13), and Pindel^47^ (v0.3). HiFi calls were produced by pbsv (v2.6.2), Sniffles^21^ (v1.0.12), cuteSV^22^ (v1.0.11), and SVision^23^ (v1.3.6). Apart from read-alignment strategies, we also used five HRAs to discover SVs, and SVs supported by at least three assemblies were included in the HRA callset (Fig. S9).

We finally obtained 9,678 large deletions and 15,324 insertions for the monozygotic twins (**Fig. 3A**). HRAs account for 92.6% of deletions and 89.3% of insertions, while HiFi reads contributed 77.1% of deletions and 68.7% of insertions, and Illumina calls covered 38.3% of deletions and 10.2% of insertions. We found that 79.8% of deletions and 75.9% of insertions could be independently supported by ONT reads (**Fig 3B**). The SV length distribution displayed ~ 300bp and ~ 6kb peaks related to SINE-Alu and LINE elements, respectively, suggesting the effective SV detection of our benchmark (**Fig. 3C, 3D,** Fig. S14). Like small variants, we also reported more high-quality variants for the monozygotic twin daughters compared to HG002 in GIAB due to the contributions of HRAs. HiFi reads and HRAs identified more SVs in repeat regions like VNTR, simple repeat, and segmental duplication regions, and variants in these complex regions were always difficult to resolve by ONT reads (**Fig. 3E, 3F**, Fig. S15, and S16). Meanwhile, SVs supported by at least two platforms always achieved a higher ONT-supporting rate compared to those supported only by one platform (**Fig 3E**). For example, there were 1,985 deletions and 4,309 insertions specifically contributed by HRAs, but around 36.0% of those calls were supported by ONT reads. Notably, 91.0% and 85.7% of HRA-specific deletions and insertions, respectively, were located in repeat regions. For example, HRAs identified a 27 kb maternal deletion at segmental duplications in *HEATR4,*but this deletion was not reported in HiFi and Illumina read alignment-based callsets (**Fig. 3G**).

**Figure 3.**
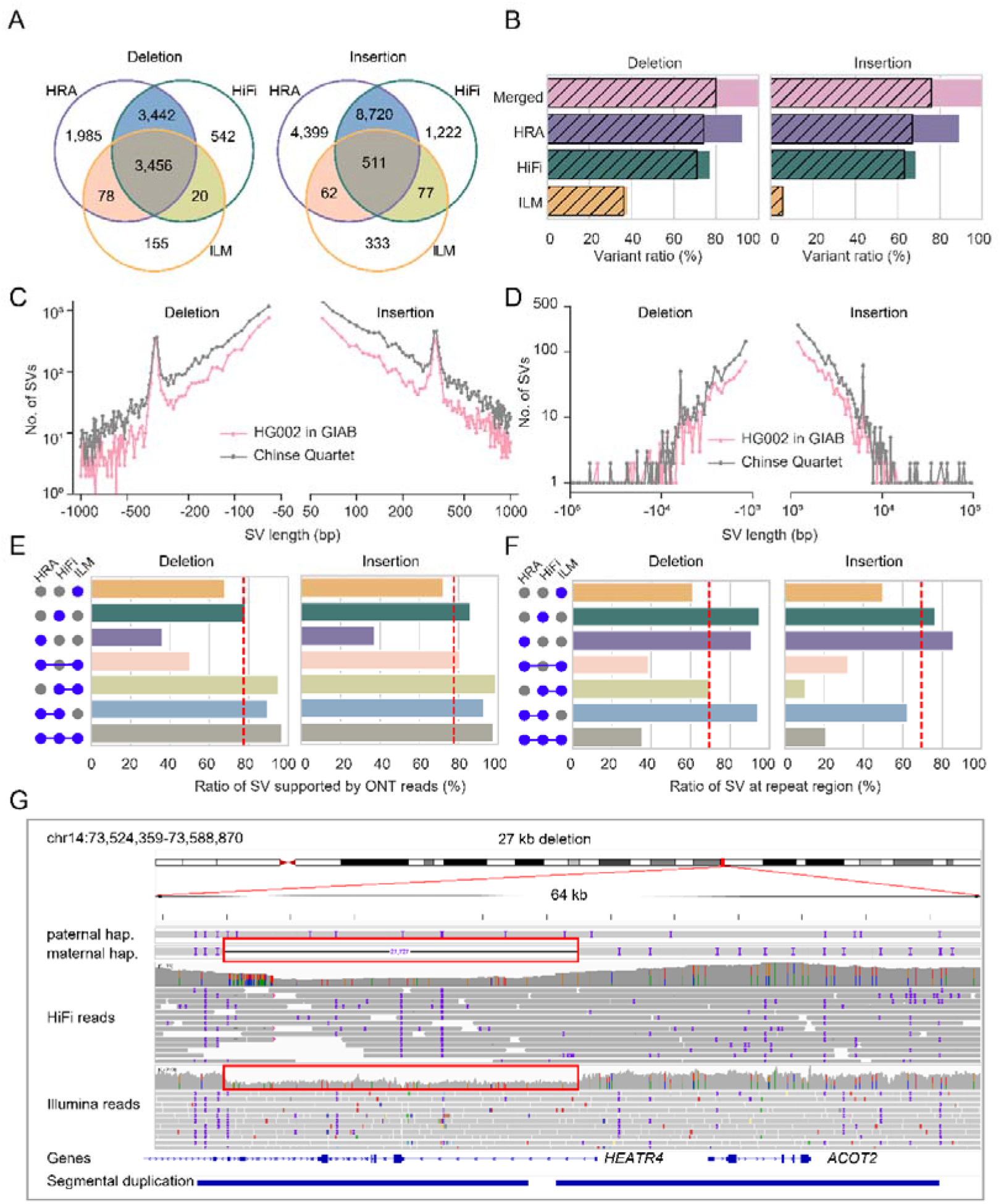
Simple SV benchmark of the Chinese Quartet. **A** Overlap of large deletions and insertions among ILM, HiFi, and HRA, respectively. **B** Bar plot depicts the percentage of ILM, HiFi, and HRA calls in the final simple SV benchmark, with gray stripes representing the supported percentages by ONT read. **C** and **D** Length distribution of large deletions and insertions in Chinese Quartet and HG002. **E** Bar plots show the rate of variation supported by ONT reads in different combinations of three technologies. **F** Bar plots represent the ratio of Indels at STR regions in different combinations of three technologies. **G** IGV snapshot shows a 27 kb deletion at a segmental duplication region.

#### Complex structural variant (CSV) and inversion benchmark construction

Detection of complex SVs and inversions was more complicated than simple variants due to ambiguous alignments, especially in repetitive regions. To build a benchmark for complex structural variants, we generated five callsets of complex SVs and inversions with HiFi reads and HRAs as input using Sniffles, SVision, cuteSV, pbsv, and PAV. Next, 175 candidate variants from the merged callset were manually inspected and refined according to IGV snapshots and dotplots (**Fig. 4A**).

**Figure 4.**
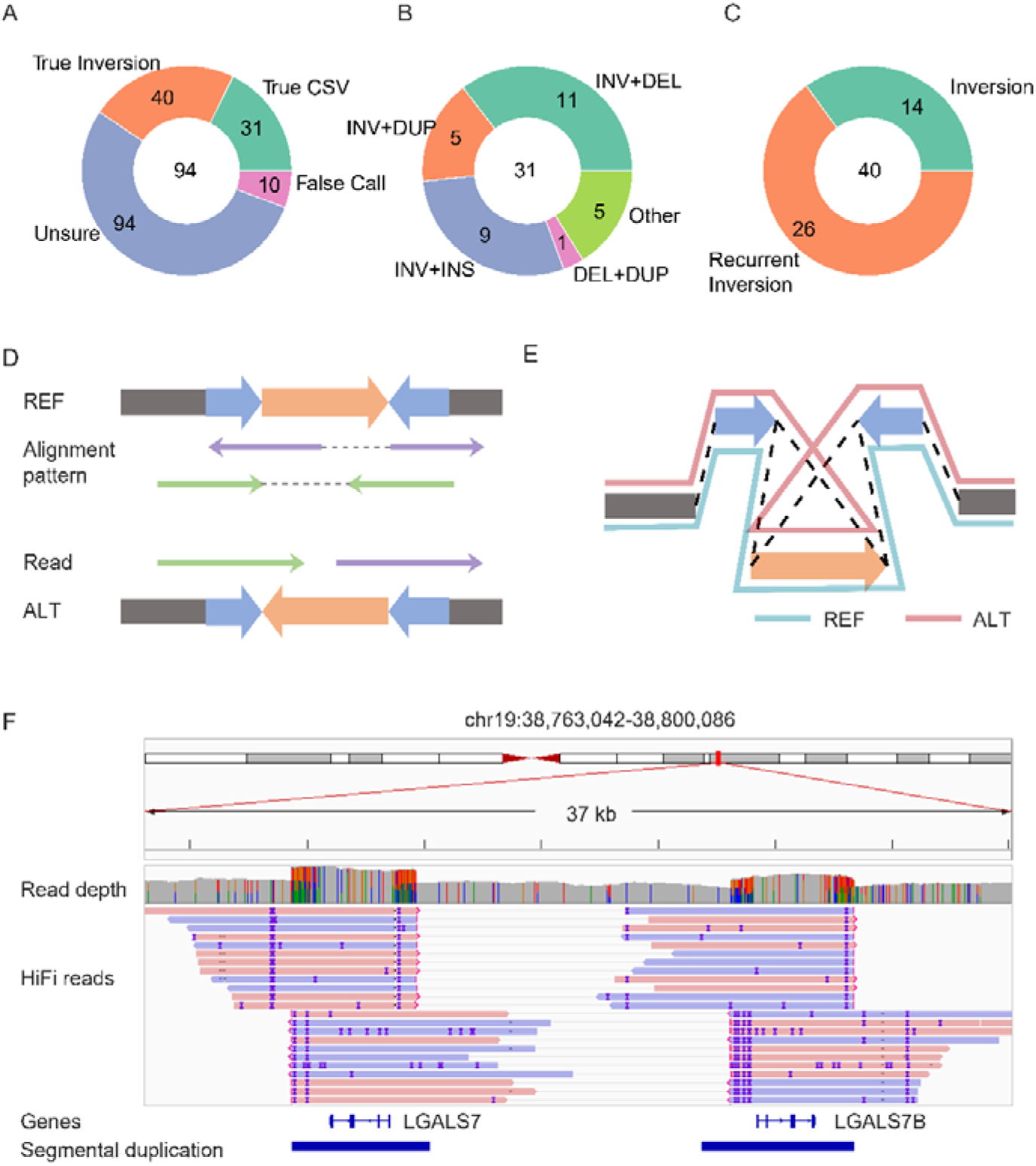
Complex SV and inversion benchmark of Chinese Quartet. **A** Composition of complex SVs and inversions. **B** and **C** The pie plot shows the composition of different types of complex SVs (B) and inversions (C) in our benchmark. **D** and **E** The diagram shows the read alignment pattern (D) and assemblies (E) of recurrent inversion. **F** The example of recurrent inversion.

Finally, we released 31 CSVs, of which 90.3% are inversion-associated (**Fig. 4B**, and Table S8). We found that Sniffles, SVision, and cuteSV discovered 80.6%-87.1% of CSVs, while PAV only reported 32.3% (Fig. S17). Only five CSVs were discovered by all callers, suggesting the challenge of CSV detection. As for inversions, we reported 40 nonredundant inversions and 75% of them were major alleles (allele frequency > 0.5) in the HGSVC callset (Table S8). We observed that 65% (26) of inversions were flanked by inverted repetitive sequences, which were defined as recurrent inversions^48^ (**Fig. 4C-F**). Notably, 92.3% of recurrent inversions were major alleles in HGSVC callset, indicating that most of recurrent inversions were caused by mis-assembly of the reference genome in such complex regions (**Fig. 4D-F**).

#### Summary and evaluation of variant benchmark

Variants in our benchmark were enriched (*P* < 1.1 × 10^−6^) in the proximal telomere of metacentric chromosomes instead of random distribution in the genome (Fig. S18 and S19). Meanwhile, the densities of SNVs and Indels are strongly correlated with the density of STR (SNV: R = 0.73, *P* = 8.35 × 10^−51^; Indel: R = 0.88, *P* = 2.58 × 10^−102^), while the densities of large deletions and insertions are strongly correlated with the density of VNTR (Deletion: R = 0.82, *P*= 5.35 × 10^−74^; Insertion: R = 0.85, *P* = 9.84 × 10^−84^) (Fig. S20). In our benchmark, we found that 27,506 SNVs, 1,003 Indels, 64 deletions, and 77 insertions affected coding DNA sequence (CDS) regions (Table S9).

In variant detection pipelines, complex regions like SD, SR, VNTR, and STR usually result in sequencing errors and multiple read alignments, particularly in short read sequencing^49^. Long read length and high base precision of HiFi and HRAs facilitated the detection of variants in complex regions, that were not accessible for other technologies (Fig. S21 and S22). Therefore, variants in our benchmark were divided into high-confidence and technology-specific calllsets according to their supporting technologies (**Fig. 5A**). In particular, variants detected by at least two technologies or also observed by either BGI or ONT reads were labeled as high-confidence calls, and variants supported only by one technology were defined as technology-specific calls. In our benchmark, technology-specific calls account for 4.4% of SNVs, 4.8 % of Indels, 14.9% of deletions, and 19.7 % of insertions. As expected, in three technology-specific callsets, 87.0% of SNVs, 94.0 % of Indels, 89.7% of deletions, and 83.0% of insertions were in repeat regions. Compared to high-confidence calls, we found that technology-specific calls always had abnormal read depths and low mappabilities due to the repetitive regions (**Fig. 5B, 5C**, and Fig. S23).

**Figure 5.**
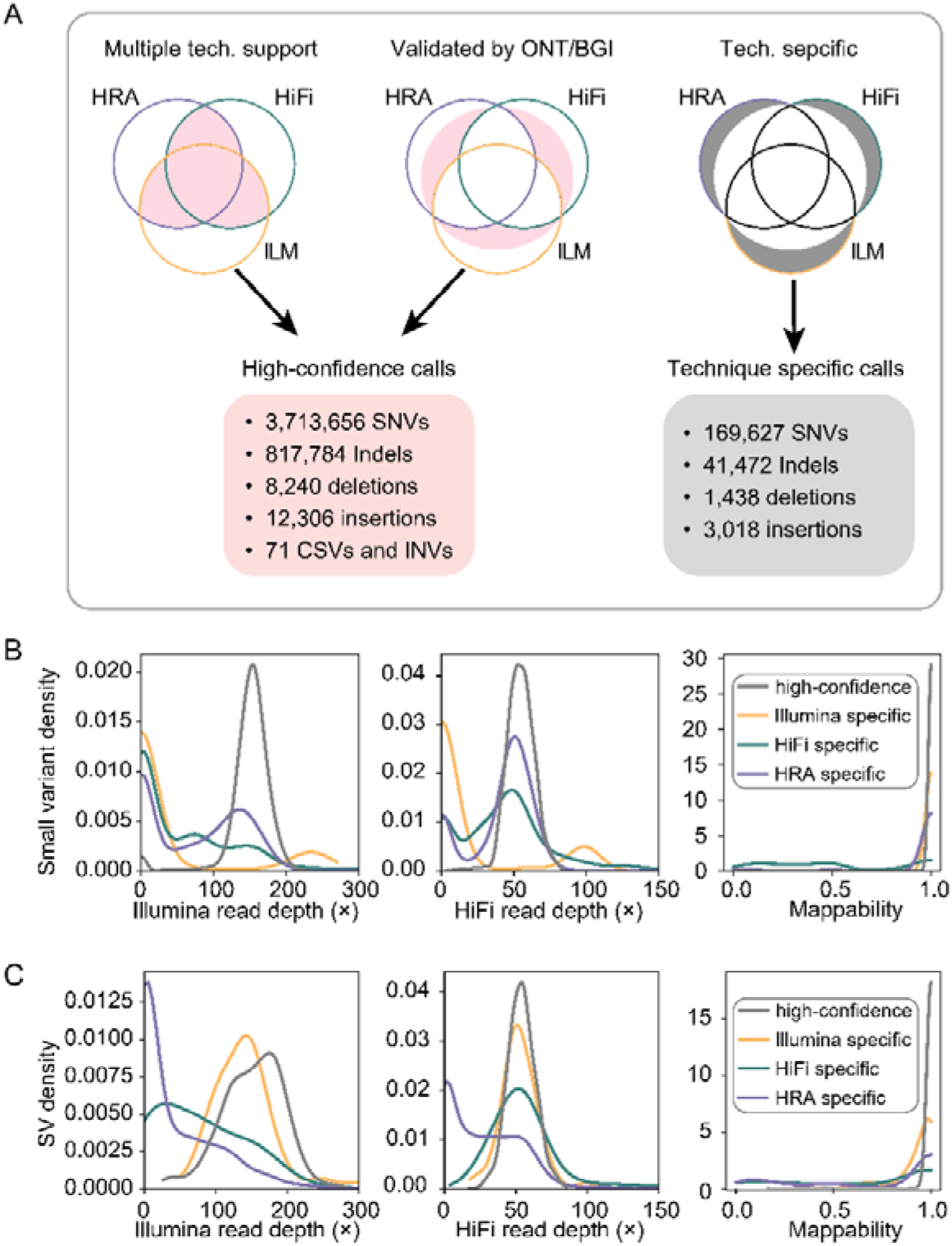
Summary and characteristics of variant benchmark. **A** Summary of variant benchmark in Chinese Quartet. **B** and **C** The density plots show the difference of variant characteristics between high-confidence and technology-specific calls in small variants (B) and structural variants (C).

### Assemblies and variant detection in different sequencing depths

Sequencing depth was an important factor for both assembly and variant detection. To further assess the assembly and variant detection pipeline in different sequencing depths, samples with multiple sequencing depths (ranging from10 × to 100 ×) were generated by downsampling the HiFi reads of monozygotic twins. Initially, samples with different sequencing depths were assembled into haplotype-resolved assemblies by hifiasm^29^. The contig N50 of two haplotypes flattened out with increasing sequencing depth and was maintained for more than 25M at 40 × (**Fig. 6A** and Table S10). The BUSCO completeness also increased rapidly and reached around 94% at 30 × (**Fig. 6A**). The accuracy of assemblies (QV) also increased steadily with the depth increase and remained stable from 60× (**Fig. 6A**, Table S10). To further evaluate the performance of variant detection with HRA in diverse sequencing depths, two haplotypes from different depths were used for variant detection with PAV^20^. Like the performance of assemblies, the recall, precision, and F1 score of variants were also improved with the increases in depth and reached a plateau at 30 × (**Fig. 6B** and Table S11). Taken together, these results suggest that 30 × HiFi reads could achieve outstanding performances in both assembly and germline variant detection pipelines.

**Figure 6.**
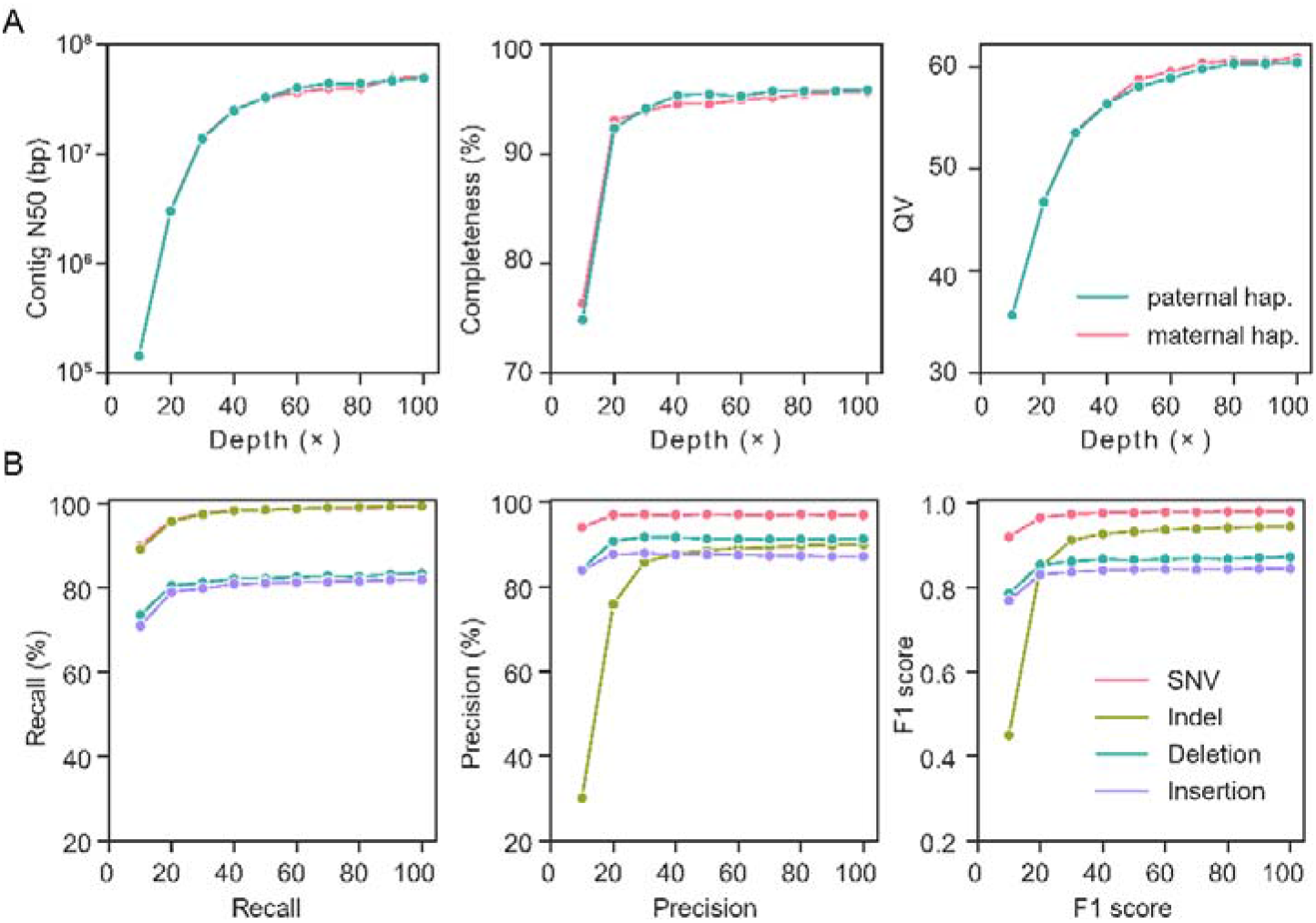
Performance of Chinese Quartet assemblies and variants of in diverse sequencing depths. **A** Contig N50 (left), completeness (middle) and QV (right) for paternal and maternal haplotypes across 10 × to 100 × sequencing depths. Completeness and QV are calculated by BUCSO and Merqury, respectively. **B** Recall, precision, and F1-score for SNVs, indels, large deletions, and insertions using assemblies with diverse sequencing depths (ranging from 10 × to 100 ×).

### Decoding HLA regions with high quality assemblies and variant benchmark

Human leukocyte antigen (HLA) genes are important in cancer, autoimmune disease, infectious disease, and tissue transplantation^50^. To better understand the genetic features of human leukocyte antigen genes, we investigated the extended major histocompatibility complex^51^ (xMHC) region of two twin daughters based on the haplotype-resolved assemblies and high-quality variant benchmark. We observed that both CQ-P and CQ-M covered the entire xMHC region in GRCh38 **(Fig. 7A**). In addition, 265 out of 271 protein-coding genes located at xMHC regions were resolved by both CQ-P and CQ-M. Compared to classical class III regions, classical class I and II had higher variant rates and lower methylation density, indicating classical class I and II regions are more active (**Fig. 7B**). We also discovered obvious distinctions in variants and methylations between two haplotypes (**Fig. 7B and 7C**). Furthermore, we discovered that the heterozygous SNVs and Indels in xMHC regions were significantly (*P* < 0.0018) more prevalent than those in other regions, while homozygous variants had no significant (*P* > 0.88) difference (**Fig. 7D**), confirming the linkage disequilibrium of HLA regions ^52^.

**Figure 7.**
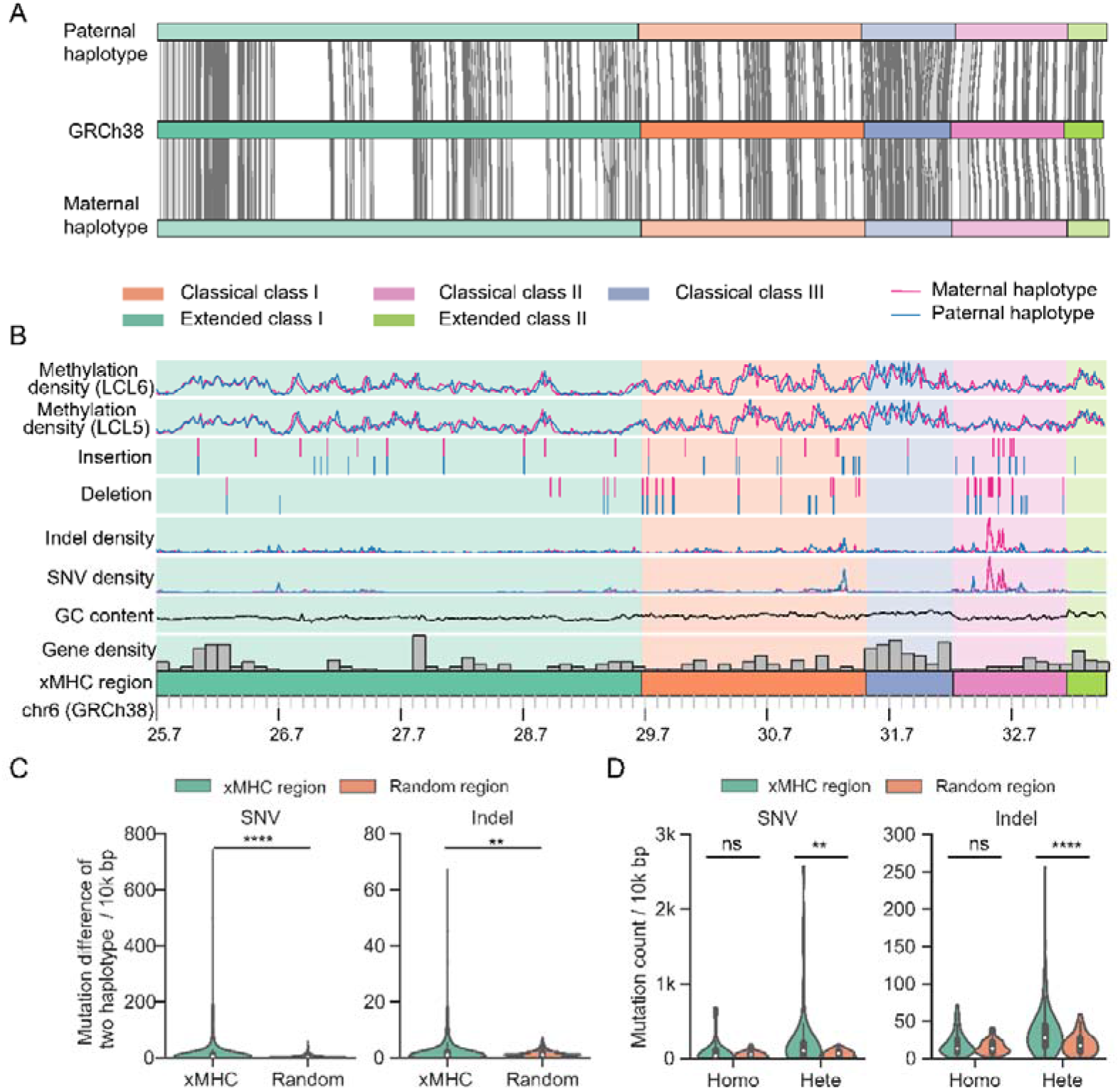
Assemblies and variants of the Chinese Quartet at extended major histocompatibility complex region. **A** Alignment of paternal and maternal haplotypes to GRCh38 at extended major histocompatibility complex (xMHC) region (chr6: 25,701,783-33,480,577). Both haplotypes covered the xMHC region with only one contigs. Gray links between haplotypes and GRCh38 are the protein coding genes resolved. **B** Genetic and epigenetic characteristics of two haplotypes. **C** Violin plot shows the variants difference between two haplotypes in 10k bp windows. The variant difference in xMHC region are significantly higher than that in other random regions (Wilcoxon rank-sum test; SNV, *P* < 0.0001; Indel *P* < 0.01). **D** Violin plot shows the heterozygous and homozygous variants count in 10k bp windows. The number of heterozygous SNVs and Indels in xMHC regions are significantly more than those in other random regions, while homozygous variants have no significant difference. ns, not significant; *, *P* < 0.05; **, *P* < 0.01; ***, *P* < 0.001; ****, *P* < 0.0001.

## Discussions

As the reference materials, the twin daughters of the Chinese Quartet could be regarded as two biological replicates, which facilitates additional cross validation than variant calling in a single sample or even in a trio. To accurately decode the reference materials, high coverage reads were generated by diverse technologies including Illumina, BGI, HiFi, and ONT. Based on the ingenious samples, the advanced data and approaches, we released high-quality haplotype-resolved assemblies for the Chinese Quartet children and constructed a comprehensive variant benchmark.

Compared to the complete hydatidiform mole (CHM13), it is more challenging to decode the complete genome of a diploid sample. Nevertheless, 76% of the chromosome arms in our assemblies of the monozygotic twins were represented by single contigs (Table S5). Meanwhile, seven and nine chromosomes of CQ-P and CQ-M were assembled at telomere-to-telomere levels, respectively (Table S5). Although advanced technologies, including HiFi and ultra-long ONT reads, were applied in our assemblies, it was still difficult to distinguish two haplotypes of diploid samples in large repetitive regions, such as higher-order repeats in centromeres. To obtain high quality assembly in these large repetitive regions, we divided the unphased reads into two haplotypes equally in our assembly pipeline. Hence, the sequences of large repetitive regions also need to be further validated by more accurate and longer reads in the future.

When including haplotype-resolved assemblies for benchmarking, more large-scale variants were detected due to the longer spanning length of HRAs on the genome^20^. Meanwhile, many variants in complex regions such as xMHC and segmental duplications were reported, which are difficult for the read-alignment strategies. Another contribution of our benchmark is that we extend the variant types to complex structural variants, compared to previous studies^7, 8, 11, 53^. Nevertheless, our benchmark also has several limitations. Firstly, technology-specific variants were subjected to further validation in the future because it was difficult for current technologies to decode all complex regions unbiasedly. For example, it is difficult for HiFi reads to resolve the variants located at large segmental duplications (Fig. 3G). Secondly, the same structural variant in our benchmark may be reported as multiple records at repeat regions due to the breakpoint shifts.

For the next phase of Chinse Quartet, we will develop new algorithms and generate novel data to improve both *de novo* assemblies and variant benchmark to facilitate resequencing projects of the Chinese Han population. We believe that the investigation of certified reference materials for genomics and other omics will prompt the reproductivity and repeatability of bioinformatics analysis in the future.

### Conclusions

In summary, we provide the high-quality haplotype-resolved assemblies and comprehensive variant benchmark for monozygotic twin daughters of the Chinese Quartet, the reference materials for whole genome-variant assessment. The high-quality assemblies and variant benchmark could be used to evaluate the performance of analysis pipelines and sequencing technologies in different centers and laboratories. For better usability of our research work, we also provide the reference materials, assemblies, and variant benchmark to the research community for improving the reproducibility of pipelines.

## Methods

### Sequencing data generation

The “Chinese Quartet” family, including father (LCL7), mother (LCL8), and two monozygotic twin daughters (LCL5 and LCL6) in this study, was from the Fudan Taizhou cohort, which was approved as certified reference material by the State Administration for Market Regulation in China. The processes of cell line establishment, DNA extraction, and Illumina sequencing were described in prior studies^12, 13^. The four cell lines were also sequenced by BGI, PacBio, and ONT solutions. Details of library preparation and sequencing in this study are described in the supplementary notes file.

### Separation of reads by haplotype

To build haplotype-resolved assemblies for the monozygotic twins of the Chinese Quartet, we split HiFi and ONT reads into paternal (CQ-P) and maternal (CQ-M) haplotypes. Firstly, we obtained the high-quality single nucleotide variants (SNVs) and Indels of the family from a previous study^13^. The variants of the monozygotic twin daughters were phased using whatshap^26^ (v1.1) with parent-child information and children’s HiFi reads. Then, we aligned HiFi, ONT, and ultra-long ONT reads of the twins to GRCh38 with minimap2^42^ (v2.20-r1061) and separated the reads into two haplotypes according to the heterozygous variants. The reads that were not covered by heterozygous variants were also assigned to the two haplotypes randomly.

### Assemblies of the Chinese Quartet

As monozygotic twins are in general regarded as genetically identical with limited somatic mutations^25^, we merged the data of two twin samples and endeavored to obtain high-quality haplotype-resolved genomes. For each haplotype of the monozygotic twin daughters, we assembled phased HiFi reads using three popular assemblers, including hifiasm^29^ (v0.15.5), hicanu^30^ (v-r10117), and flye^28^ (v2.8.3-b1695). Meanwhile, ONT regular and ONT ultra-long reads were assembled with flye^28^ (v2.8.3-b1695) and shasta^27^ (0.7.0). Next, we identified the mis-assembly and broke the chimeric contigs with ragtag^31, 54^ (v2.0.1). Then we scaffolded the hifiasm contigs based on the human Telomere-to-Telomere genome^19^ (CHM13 v1.0) and closed the gaps of hifiasm scaffolds with other contigs by Gapless (https://github.com/PengJia6/gapless). Finally, two haplotypes were polished with corresponding HiFi reads using NextPolish^32^ (v1.3.1).

### Assemblies evaluation and analysis

Two haplotype-resolved assemblies of the monozygotic twin daughters were evaluated in three aspects, including accuracy, continuity, and completeness. The accuracies (QV score) of the Chinese Quartet genomes were evaluated according to the Illumina reads by Merqury^55^ (v1.3). For continuity evaluation, we calculated contig numbers, contig N50, and the gap of HRAs. In terms of completeness, we applied three methods to evaluate CQ-P and CQ-M. First, we applied BUSCO^39^ (v5.1.3) with mammalia_odb10 to calculate the fraction of complete BUSCO genes. Then, Merqury^55^ (v1.3) was used to estimate the completeness of HRAs with Illumina sequencing data. Meanwhile, we aligned our assemblies to GRCh38 with minimap2, and the coverage fractions of our assemblies to GRCh38 were calculated for completeness assessment.

### Novel sequences identification and genome annotation

We aligned contigs of two haplotypes of the Chinese Quartet to GRCh38 with minimap2^42^ (v2.20-r1061) and winnowmap2^56^ (v2.03). Thereafter, the sequences labeled by hard-clip (H), soft-clip (S), and insertion (I) in bam files were extracted and aligned to GRCh38 again. The unmapped sequences were collected as novel sequences. We also annotated the protein-coding genes of our assemblies by Liftoff^40^ (v1.6.1) based on gencode annotation (v38) of GRCh38. Then, the novel sequences of our assemblies were extracted and repeat regions were marked by RepeatMasker (v4.1.2-p1, http://www.repeatmasker.org). Finally, the unannotated regions were further annotated by Augustus^41^.

### Variant detection of the Chinese Quartet by Illumina reads

The SNVs and Indels of the Chinese Quartet by Illumina were downloaded from a previous study^13^. To discover structural variants using short reads, we aligned Illumina reads to GRCh38 and marked duplication reads with biobambam2^57^ (v2.0.182). Then we detected variants of the Chinese Quartet by Manta^44^ (v1.6.0), Delly^45^ (v0.9.1), Lumpy^46^ (v0.2.13), and Pindel^47^ (v0.3). We kept the SVs with at least 30 reads supporting and 50 bp long for the following steps. Only SVs supported by both girls and one of their parents were kept as high-quality variants of the twins for each caller. High quality variants from four callers were then integrated by Jasmine^58^ (v1.1.5) for each SV type, respectively. Finally, variants supported by at least two callers were retained for the final benchmark.

### Variant detection of the Chinese Quartet by HiFi reads

We aligned HiFi reads to GRCh38 using minimap2^42^ (v2.20-r1061) and then detected small variants for each sample using deepvariant^43^ (v1.1.0) with the parameter “--model__type=PACBIO” set. The GVCFs of four samples were merged and genotyped by glnexus (v1.2.7). SNVs and Indels were phased according to parent-child information and children’s HiFi reads^26^ (v1.1). To obtain high-quality SNVs and Indels, we filtered variants in four steps: (i) filtering variants with allele frequencies less than 0.2, read depth less than 25 or more than 75; (ii) removing variants violating the Mendelian rule, (iii) only keeping variants, of which two girls had same genotypes; (iv) filtering variants longer than 49bp.

To obtain high-quality SV calls from the Chinese Quartet, we utilized four popular callers, including pbsv (v2.6.2), Sniffles^21^ (v1.0.12), cuteSV^22^ (v1.0.11), and SVision^23^ (v1.3.6), to discover SV events. Similar to Illumina reads, we also kept SVs with at least 15 reads supported. Then SVs following the Mendelian rule and supported by at least two callers were kept for the final benchmark.

### Variant detection of the Chinese Quartet by HRAs

Apart from read-alignment strategies, contigs of HARs were also used for variant detection. We aligned HRAs to GRCh38 using minimap2^42^ (v2.20-r1061) and discovered variants by PAV^20^ (v1.1.0) pipelines. We discovered SNVs and Indels with HiFi assemblies, and only variants detected by all three assemblies were kept in the final benchmark. SVs were discovered by both HiFi and ONT assemblies, and we kept variants with at least two assemblies supported in the final benchmark.

### Complex structural variant and inversion detection

To expand the complex structural variants in our benchmark, we used HiFi reads and HRA to discover variants. In raw callsets, SVs labeled by multiple types, inversion, and CSV were extracted as candidate variants. All candidate variants were manually refined by IGV snapshots and dotplots. In particular, variant types were determined by the dotplots between HRAs and the reference genome.

### Variant benchmark construction and evaluation

SNVs and Indels calls from Illumina, HiFi, and HRAs were merged with bcftools (v1.13) and large deletions and insertions were merged with Jasmine^58^ (v1.1.5). Variants at centromeres, telomeres, copy number abnormal regions, and sex chromosomes were excluded in the final benchmark. To evaluate the quality of SNVs and Indels in our benchmark, BGI reads were aligned to GRCh38, and the deepvariant^43^ was used to call SNVs and Indels. ONT reads were aligned to the reference genome and four callers, including pbsv (v2.6.2), Sniffles^21^ (v1.0.12), cuteSV^22^ (v1.0.11), and SVision^23^ (v1.3.6), were used to call variants. We kept SVs supported by at least 15 reads and 2 callers in ONT for SV evaluation. In our benchmark, variants that were supported by at least two technologies or supported by BGI or ONT reads were labeled as “high-confidence” calls. Moreover, the variants only detected by one technology were assigned as “technology-specific” calls.

### Chinese Quartet benchmark annotation

Repeat regions, including segmental duplication (SD), simple repeat (SR), variable number tandem repeat (VNTR), and repeat mask (RM), were downloaded from the table browser. Short tandem repeats (STRs) were generated by the “scan” command in msisensor-pro^59^. A variant was annotated to repeat regions if it overlapped with repeat regions. Variants were also annotated by the Ensembl Variant Effect Predictor (VEP) ^60^ (v104.3).

### Haplotype-resolved methylation calling of Chinese Quartet

Phased ONT reads and raw fast5 files were indexed with the “index” command of nanopolish^61^ (0.13.2). Then we call methylation of each haplotype using phased reads with the “call-methylation” command in nanopolish. Next, the methylation frequency of each site was calculated by the ‘calculate_methylation_frequency.py’. Finally, we kept the sites with a methylation frequency greater than 0.8 as methylated sites.

### Chinese Quartet benchmark application

To further assess the performance of assemblies and variants calling in various sequencing depths, we downsampled the HiFi reads of two monozygotic twins ranging from 10 × to 100 with increments by 10 ×. We assembled the simulated samples from different sequencing depths with hifiasm^29^ and called variants by PAV^20^ pipelines. The assemblies were evaluated in three aspects, including accuracy, completeness, and continuity, as in the previous description. As for variants, only calls supported by both LRA and minimap2 in the PAV pipeline remained as high-quality calls. We defined the variant supported by both the benchmark and simulated sample as “true positive” (TP) call. The variant only supported by the simulated sample and benchmark was labeled as “false positive” (FP) and “false negative” (FN) call, respectively. Then, recall, precision, and F1 score of variant detection were calculated by equations (1–3).

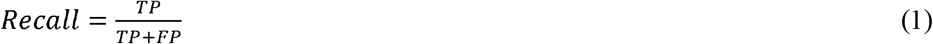

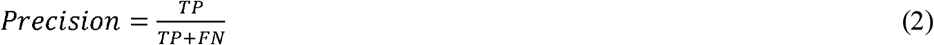

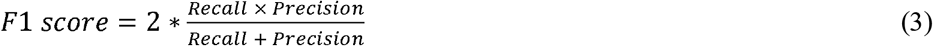

## Authors’ contributions

Conceptualization: K.Y., J.W., L.S., P.J., and L.D. Sequencing data generation: L.D., YUA.Z., YUJ.Z., X.W., F.L., and Y.W. Data management and archiving: P.J., L.D, YUA. Z., and L.R. Genome assembly: P.J., K.Y., B.W., X.Y., X.Z, and J.R. Variant analysis: P.J., K.Y., J.L., T.W., and S.W. Software and pipeline development: P. J. Validation: P.J. and L.R. Visualization: P.J., T.X., N.D., and Y.C. Organization of supplementary materials: P.J. Original manuscript writing: P.J. and K.Y. Manuscript review and editing: K.Y., P.J., B.W., X.Y. and L.D. Project administration and supervision: K.Y. and J.W.

## Availability of data and materials

The Certified Reference Materials can be requested from the Quartet Data Portal (http://chinese-quartet.org/) under the Administrative Regulations of the People’s Republic of China on Human Genetic Resources. All raw sequencing reads of the reference materials have been deposited in the Genome Sequence Archive^62^ at the National Genomics Data Center, Beijing Institute of Genomics, Chinese Academy of Sciences/China National Center for Bioinformation (GSA: HRA001859), and are publicly accessible at https://ngdc.cncb.ac.cn/gsa. The assemblies and variant benchmark are also available from GSA (PRJCA007703) or from the authors upon request. Other supporting data is available at the additional files of this paper or from the authors upon request. Pipelines for genome assembly and variant detection are available at Github (https://github.com/xjtu-omics/ChineseQuartetGenome).

## Acknowledgments

We would like to thank Guangbo Tang, Zihang Li, and Xiujuan Li for the cell culturing in this project and Jing Hai and Huanhuan Zhao for administrative and technical support.

