## Supplementary notes for "Haplotype-resolved assemblies and variant benchmark of a Chinese Quartet"

### 1 Generation of sequencing data

#### 1.1 Library preparation and sequencing of HiFi reads

SMRTbell target size libraries were constructed for sequencing according to PacBio's standard protocol (Pacific Biosciences, CA, USA) using 15kb preparation solutions. The main steps for library preparation are: (1) gDNA shearing, (2) DNA damage repair, end repair and A-tailing, (3) ligation with hairpin adapters from the SMRTbell Express Template Prep Kit 2.0 (Pacific Biosciences), (4) nuclease treatment of SMRTbell library with SMRTbell Enzyme Cleanup Kit, (5) size selection, and (6) binding to polymerase. Briefly, a total amount of 15 μg DNA per sample was used for the DNA library preparations. The genomic DNA sample was sheared by gTUBEs (Covaris, USA) according to the expected size of the fragments for the library Single-strand overhangs were then removed, and DNA fragments were damage repaired, end repaired and A-tailing. Then the fragments ligated with the hairpin adaptor for PacBio sequencing. And the library was treated by nuclease with SMRTbell Enzyme Cleanup Kit and purified by AMPure PB Beads. Target fragments were screened by the BluePippin (Sage Science, USA). The SMRTbell library was then purified by AMPure PB beads, and Agilent 2100 Bioanalyzer (Agilent technologies, USA) was used to detect the size of library fragments. Sequencing was performed on a PacBio Sequel II instrument with Sequencing Primer V2 and Sequel II Binding Kit 2.0 in Grandomics.

#### 1.2 Library preparation and sequencing of regular ONT reads

A total amount of 3-4 μg DNA per sample was used as input material for the ONT library preparations. After the sample was qualified, size-select of long DNA fragments were performed using the Pippin HT system (Sage Science, USA). Next, the ends of DNA fragments were repaired, and A-ligation reaction were conducted with NEB Next Ultra II End Repair/dA-tailing Kit (Cat# E7546). The adapter in the SQK-LSK109 (Oxford Nanopore Technologies, UK) was used for further ligation reaction and DNA liabrary was measured by Qubit® 4.0 Fluorometer (Invitrogen, USA). About 700ng DNA library was constructed and performed on a Nanopore PromethION sequencer instrument (Oxford Nanopore Technologies, UK) at the Genome Center of Grandomics (Wuhan, China).

#### 1.3 Library preparation and sequencing of Ultra-long ONT reads

For each ultra-long Nanopore library, approximately 8-10 µg of gDNA was sizeselected (>50 kb) with SageHLS HMW library system (Sage Science, USA), and processed using the Ligation sequencing 1D kit (SQK-LSK109, Oxford Nanopore Technologies, UK) according the manufacturer' s instructions. About 800ng DNA libraries were constructed and sequenced on the Promethion (Oxford Nanopore Technologies, UK) at the Genome Center of Grandomics (Wuhan, China).

### 2 Details for variant benchmark establishing

#### 2.1 Small variants detection

##### 2.1.1 Small variants detection by Illumina reads

High quality SNVs and Indels of Chinese Quartet were obtained from https://zenodo.org/record/5275189#.YaaYn9DMJPZ [1]. We merged the variants of form samples with bcftools (v1.12), then phased the variants by whatshap [2] (v1.1). Variants violating the Mendelian rule were removed in the final benchmarks.

##### 2.1.2 Small variants detection by HiFi reads

HiFi reads were aligned to GRCh38 using minimap2 [3] (v2.20-r1061) with parameters "-a -H -k19 -O 5,56 -E 4,1 -A 2 -B 5 -z 400,50 -r 2000 -g 5000 --eqx --MD -Y " setting and then sorted by samtools. Subsequently, we called small variants for each sample using deepvariant [4] (v1.1.0) with flowing parameter "--model_type=PACBIO" setting. GVCFs of four samples were merged and genotyped by glnexus (v1.2.7). SNVs and

Indels were phased according to their and HiFi reads by whatshap. To obtain high-quality calls, we filtered variants with four steps: (i) filtering variants with allele frequencies less than 0.2, read depth less than 25 or more than 75, (ii) removing variants violating the Mendelian rule, (iii) only keeping variants, of which two girls had same genotypes, (iv) filtering variants longer than 49bp.

##### 2.1.3 Small variants detection by haplotype-resolved assemblies

HiFi reads of the twin daughters were applied to hifiasm, hicanu, and flye. Then, we discovered variants with three haplotype-resolved assemblies using PAV pipelines. Only variant supported by all three callsets was kept in following steps.

#### 2.2 Structural variants detection

##### 2.2.1 Structural variant detection by Illumina reads

Illumina reads of four samples were aligned to GRCh38 with bwa [5] (v0.7.17-r1188) mem command and the aligned reads were sorted by samtools [6] (v1.12). Next, PCR duplicated reads were marked by biobambam2 (v2.0.182). Then, we called variants of canonical chromosomes (chromosome 1 to 22, X and Y) and mitochondrial DNA for Chinese Quartet samples by Manta[7] (v1.6.0), Delly[8] (v0.9.1), Lumpy[9](v0.2.13), and Pindel[10] (v0.3).

For Manta, we used default parameters for SV calling, and only kept the records with PASS filters. For Delly, we used default parameters and excluded regions of the blacklist file in the HGSVC2 data portal with '-x' parameter and kept the records with PASS filters. For Lumpy, we run the lumpy by Smoove (v0.2.8) call command with. '-x' and '--genotype' parameters setting and also excluded of the blacklist file in the HGSVC2 data portal with '--exclude' parameter. For Pindel, we call variants with '-x2 -l' parameter setting and excluded the blacklist file in the HGSVC2 data portal with '-J' parameter. Then, pindel format output was transferred to vcf format by pindel2vcf (v0.6.4) with '-R GRCh38 -is 50 -as 100000000 -b -e 10 -ss 5' parameters setting.

SVs shorter than 50bp were removed and only deletions (DELs), duplications (DUPs), insertions (INSs), and inversions (INVs) of canonical chromosomes were kept for following steps. Subsequently, four filtered callsets of each sample were integrated using Jasmine [11] (v1.1.5). Then, the merged SVs of four samples were concatenated by Jasmine and only SV longer than 49bp were kept in final callset. Finally, we filtering the variants violating the Mendelian rule in benchmark callset.

##### 2.2.2 Structural variant detection by HiFi reads

To obtain high-quality SV calls from HiFi reads of the Chinese Quartet, we utilized four popular callers, including pbsv (v2.6.2), Sniffles [12] (v1.0.12), CuteSV [13] (v1.0.11), and SVision (v1.3.6), to discover SV events and integrate their calls by Jasmine. For pbsv, we called SVs with "-t DEL,INS,INV,DUP,BND,CNV --ccs " setting. For Sniffles, we detected SVs with "–s 3". For cuteSV, we discovered SVs with "--max_cluster_bias_INS 1000 --diff_ratio_merging_INS 0.9 -s 2 --genotype --diff_ratio_merging_DEL 0.5 --max_cluster_bias_DEL 1000 --min_size 20 " setting. SVision calls were generated using the "--min_sv_size 30 -s10". For each caller, variants of four samples were filtered according to the read depth, allele depth, and Mendelian rule. Finally, only variants supported by at least two callers were kept in the following analysis.

##### 2.2.3 Structural variant detection by haplotype-resolved assemblies

We assembled ONT reads using shasta and flye and HiFi reads using hifiasm, hicanu and flye, obtaining five haplotype-resolved assemblies for each haplotype. Then, we discovered structural variants with five haplotype-resolved assemblies using PAV pipelines. Only variant supported by at least three callsets was kept in following steps.

#### 2.3 Complex variants and inversion detection

To discover the complex structural variants in our benchmark, we obtained complex variants from five callsets. HiFi reads were applied to pbsv (v2.6.2), Sniffles [12] (v1.0.12), CuteSV [13] (v1.0.11), and SVision (v1.3.6) [14], and SVs labeled as Inversion, CSV, and multiple types were extracted as candidates. The inversion of HRA calls were also extracted as candidates. The sequencing alignments of Illumina reads, HiFi reads, and HRAs in candidate regions were visualized by IGV [15]. The dotplot between HRAs and the reference genome of candidate regions was generated by Gepard [16]. The candidates were further refined and validated by the IGV snapshots and dotplots.

#### 3 Variant benchmark evaluation

To assess the validated rate of each variant type in our benchmarks, we calculated the validated rate of each variant type in our benchmarks by equation (1-5).

$$P^{m,v} = \frac{N^{m,v}}{N^{m}} \quad (1)$$

$$P_t^{m,v} = \frac{N_t^{m,v}}{N^{m,v}} \quad (2)$$

$$P_t^{m,s} = \frac{N_t^{m,s}}{N^{m,s}} \quad (3)$$

$$N^{m,s} = N_{ILM}^{m,s} + N_{HiFi}^{m,s} + N_{HRA}^{m,s} \quad (4)$$

$$N^{m,v} = N^{m} - N^{m,s} \quad (5)$$

Where $m \in \{SNV,\ Indel,\ Deletion,\ Insertion\}$ and $t \in \{ILM,\ HiFi,\ HRA\}$. $P^{m,v}$ was the validated rate of *m* variant type (Fig. 2D). $N^{m}$ and $N^{m,v}$ were total count and validated count of variant type *m*, respectively. $N_t^{m,v}$ and $N_t^{m,s}$ denoted the number of validated and singleton variant detected by technology *t*. $N^{m,s}$ represented the total singleton variants number in our benchmarks. $P_t^{m,v}$ (x-axis of Fig. 2E) was used to denote the ability of technology *t* to detect validated variants. And $P_t^{m,s}$ (y-axis of Fig. 2E) was the ability of technology *t* to detect specific variants.

#### 4 Code and data availability

The detailed description of Certified Reference Materials is available at Quartet Data

Portal (http://chinese-quartet.org/). The Certified Reference Materials can be requested from the Quartet Data Portal (http://chinese-quartet.org/) under the Administrative Regulations of the People's Republic of China on Human Genetic Resources. All raw reads (FASTQ files) of the standard reference materials are available from the Genome Sequence Archive[17] at the National Genomics Data Center, Beijing Institute of Genomics, Chinese Academy of Sciences/China National Center for Bioinformation (GSA: HRA001859), and are publicly accessible at https://ngdc.cncb.ac.cn/gsa. Data description of the standard reference are available at https://docs.chinese-quartet.org/.

Variants of the standard reference by Illumina reads are downloaded from https://zenodo.org/record/5275189#.YaaYn9DMJPZ. Small variants for HG002 are downloaded from GIAB FTP site at https://ftp-trace.ncbi.nlm.nih.gov/ReferenceSamples/giab/release/AshkenazimTrio/HG002_NA24385_son/latest/GRCh38/HG002_GRCh38_1_22_v4.2.1_benchmark.vcf.gz. Large variants for HG002 are downloaded from https://ftp-trace.ncbi.nlm.nih.gov/ReferenceSamples/giab/data/AshkenazimTrio/analysis/NIST_SVs_Integration_v0.6/HG002_SVs_Tier1_v0.6.vcf.gz. Variants for HGSVC samples are downloaded at http://ftp.1000genomes.ebi.ac.uk/vol1/ftp/data_collections/HGSVC2/.

The final assemblies and variants benchmarks are available at https://stuxjtueducn-my.sharepoint.com/personal/pengjia_stu_xjtu_edu_cn/_layouts/15/onedrive.aspx?id=%2Fpersonal%2Fpengjia%5Fstu%5Fxjtu%5Fedu%5Fcn%2FDocuments%2Fsafetyshare%2FChineseQuartet&ga=1. The website of Chinese Quartet project is available at https://github.com/xjtu-omics/ChineseQuartetGenome. The pipeline for assembly is available at https://github.com/PengJia6/AssmPipe. The assembly merging pipeline is available at https://github.com/PengJia6/gapless. The pipeline for assembly evaluation is available at https://github.com/PengJia6/Postassm. The pipelines for variant detection are available at https://github.com/PengJia6/NGSGemlineMutPipe and https://github.com/PengJia6/TGSGermlineMutPipe. Other supporting data are available in the Additional files.

### 5. List of Supplementary

#### 5.1 List of supplementary tables

Table S1. Sequencing summary of the Chinese Quartet.

Table S2. Long read summary of two haplotypes for assembly.

Table S3. Assembly summary of the Chinese Quartet by different assemblers and sequencing technologies.

Table S4. Assembly summary of the Chinese Quartet and other samples.

Table S5. Distribution of contigs in Chinese Quartet assemblies.

Table S6. The aligned fraction to GRCh38 of the Chinese Quartet haplotypes.

Table S7. Annotations of novel genes in Chinese Quartet

Table S8. Summary of Complex SVs and inversion of Chinese Quartet.

Table S9. Summary of VEP annotations for benchmarks.

Table S10. Assembly performance at different sequencing depths of the Chinese Quartet.

Table S11. Variant detection performance at different sequencing depths of the Chinese Quartet.

#### 5.2 List of supplementary figures

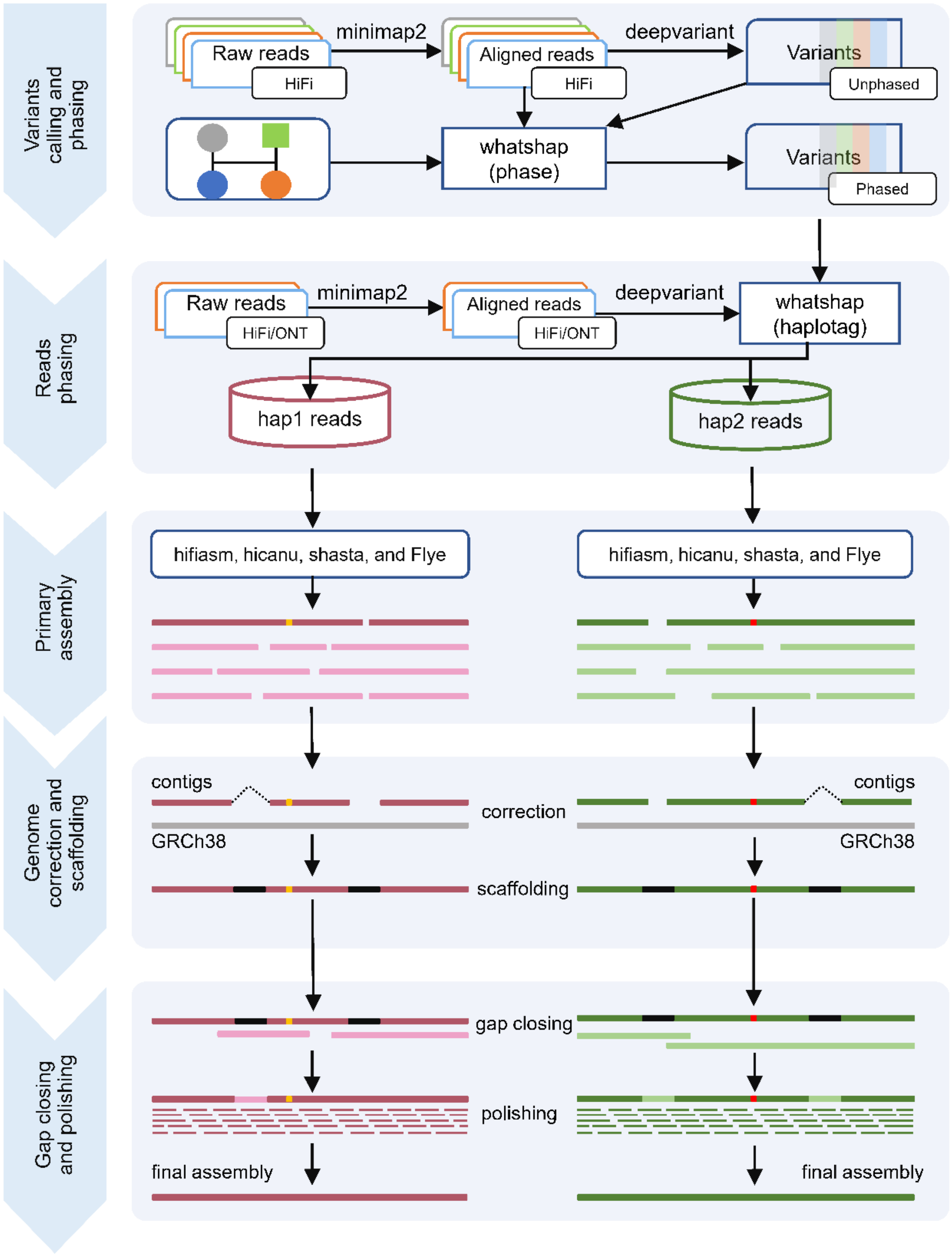

**Figure S1** Haplotype-resolved assembly pipeline for the Chinese Quartet twins.

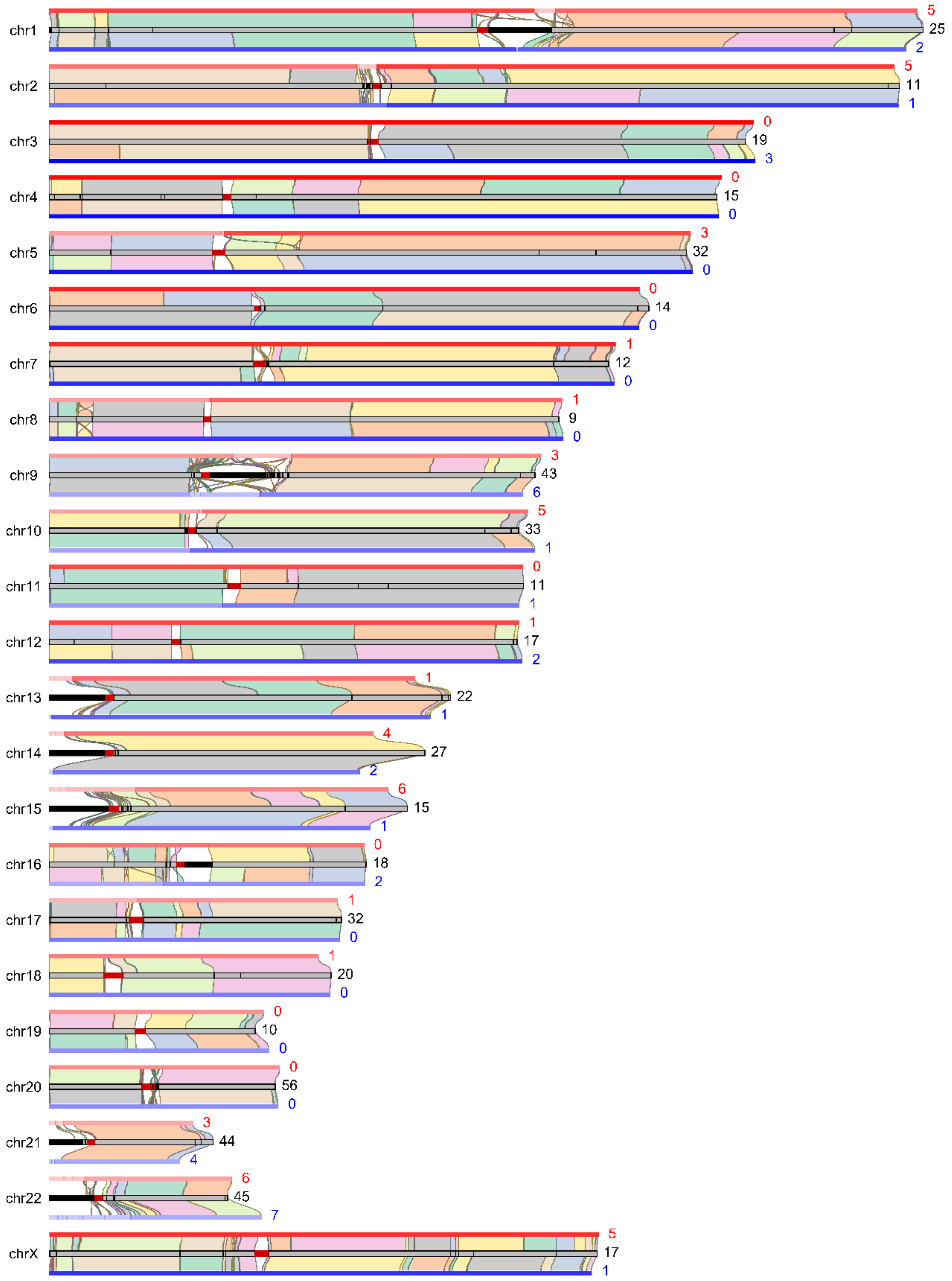

**Figure S2** Alignments of paternal (top) and maternal (bottom) haplotypes to GRCh38. The color shading of two haplotypes represents the contig size, with darker segments for longer contigs. The numbers of the gaps in CQ-P, CQ-M, and GRCh38 are labelled on the right of the chromosomes.

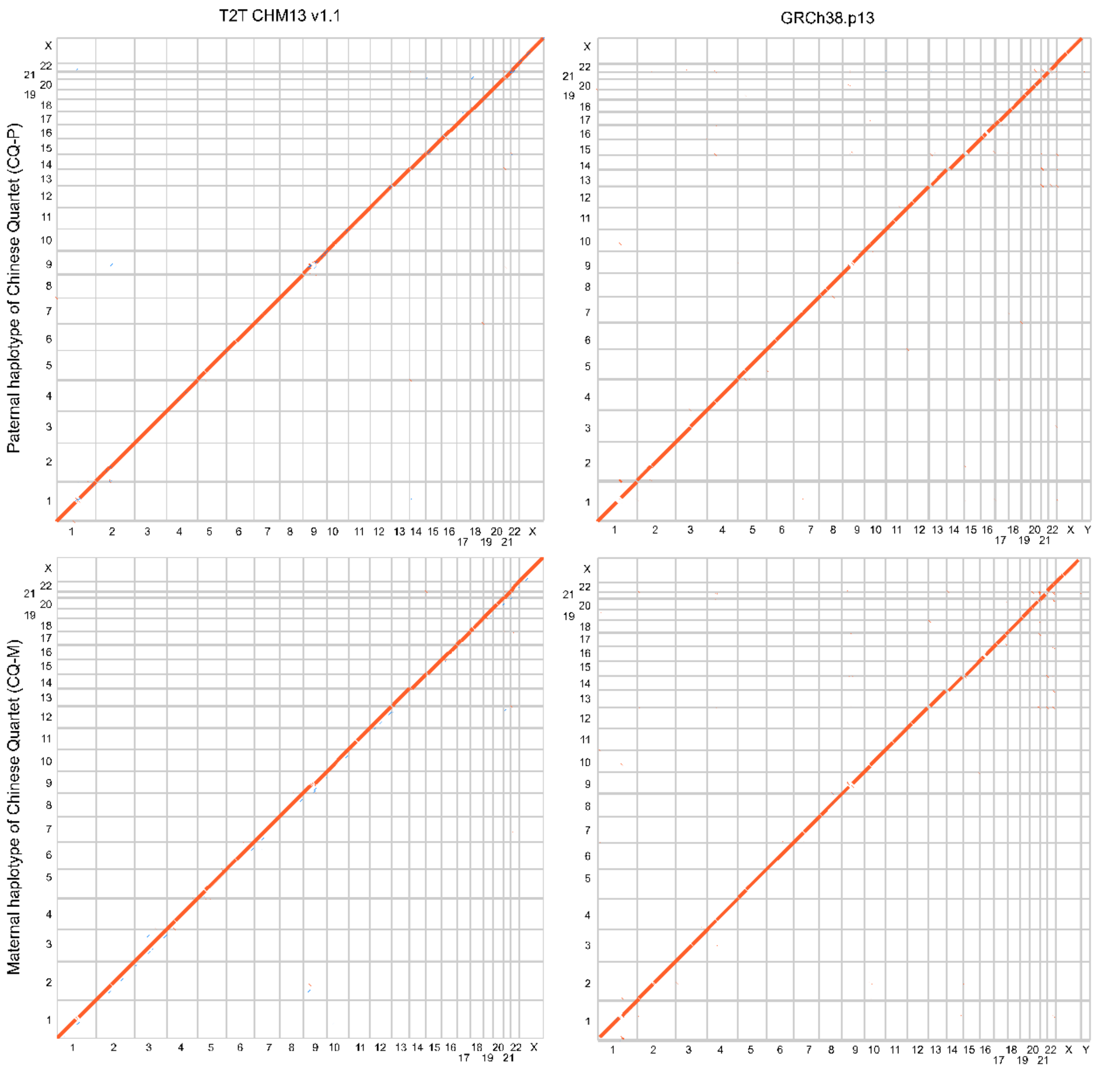

**Figure S3** Comparison between haplotype-resolved assemblies of the Chinese Quartet twin daughters and GRCh38 as well as CHM13. Dotplots represent the co-linearity between CQ-P/CQ-M and GRCh38/CHM13.

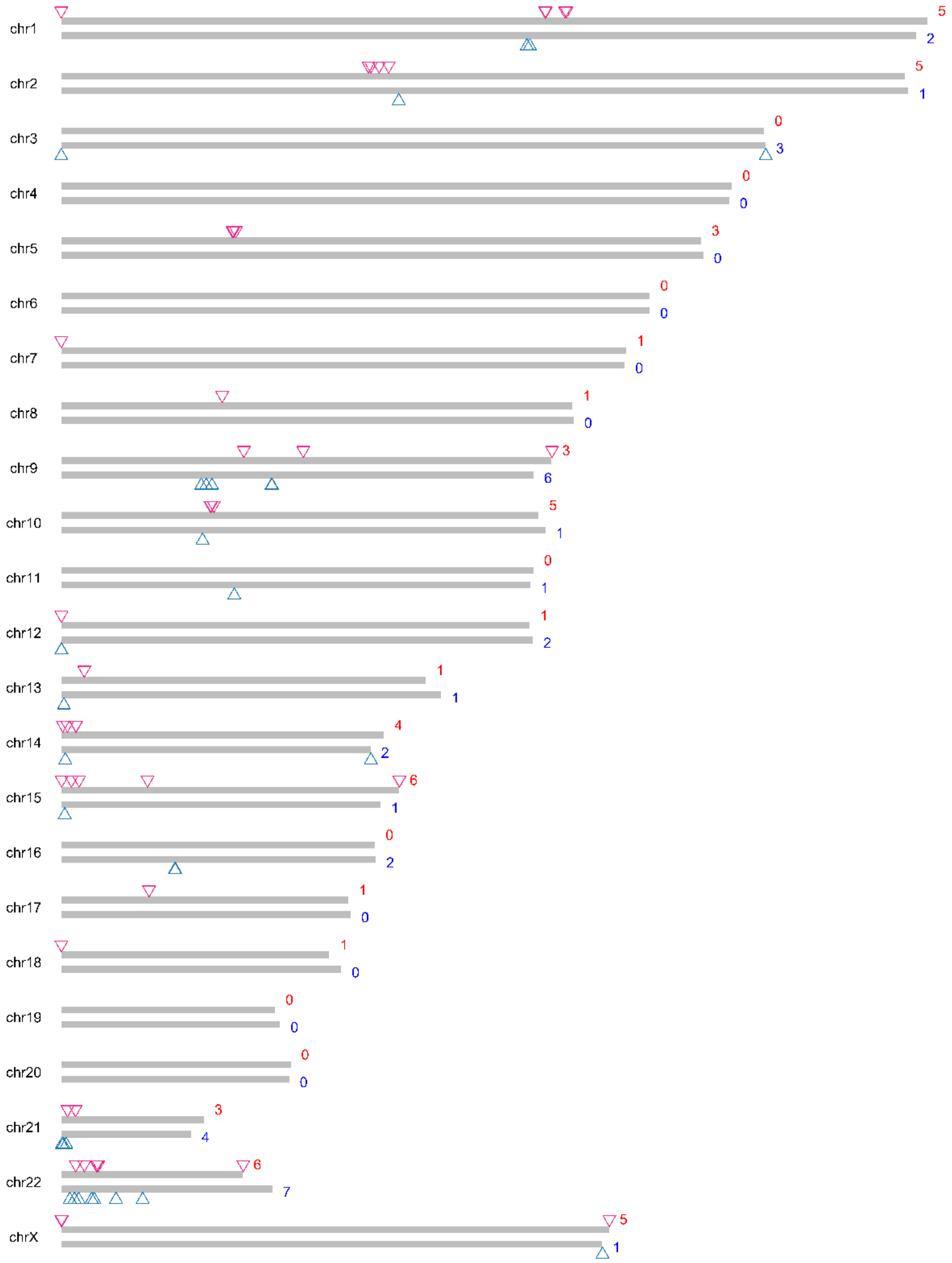

**Figure S4** Overview of the gaps in the assemblies of Chinese Quartet twin daughters. Triangles denote the locations of gaps on CQ-P (top) and CQ-M (bottom). The gap numbers of CQ-P and CQ-M are labelled on the right of the ideogram.

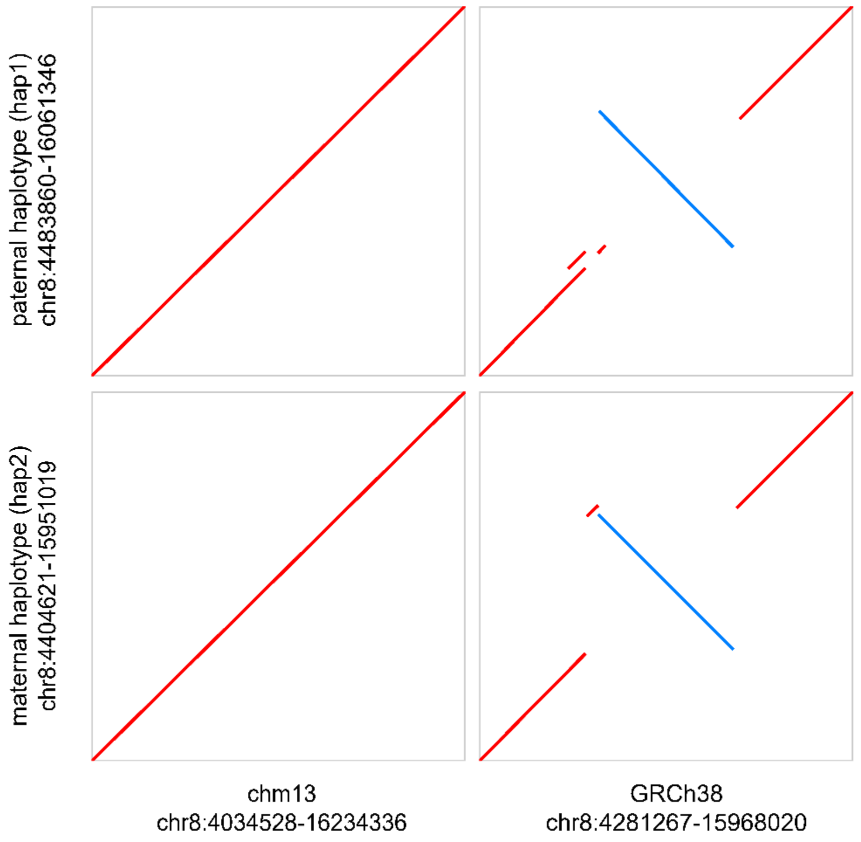

**Figure S5** Dot plots demonstrate that CQ-P and CQ-M have a ~ 4M inversion at chromosome 8 when compared to GRCh38 but are consistent with the sequences from the T2T genome.

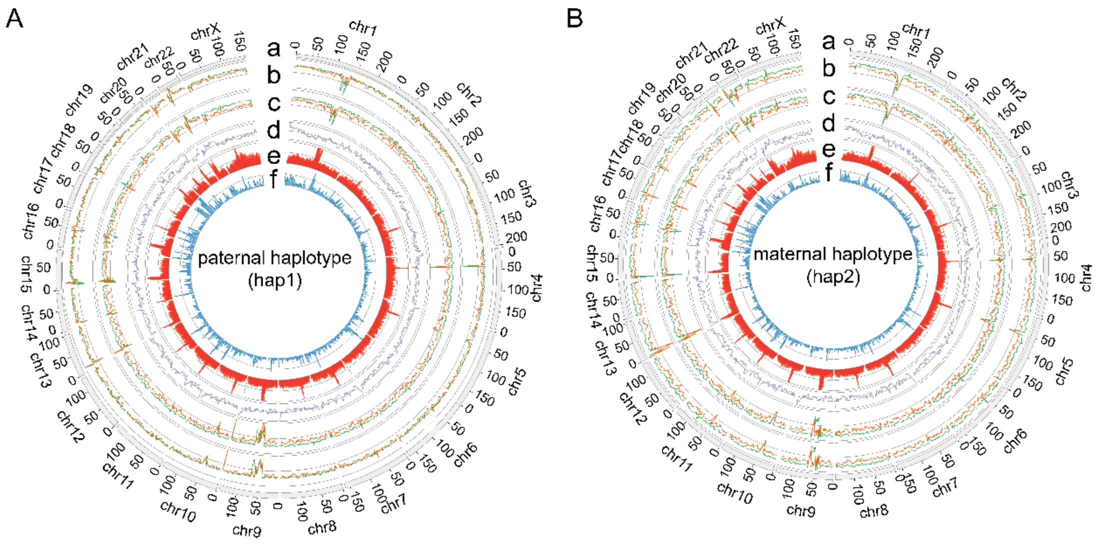

**Figure S6** Circos plots show the characteristics of the paternal (A) and maternal (B) haplotypes of the Chinese Quartet twin daughters. Track a denotes the ideogram. Track b and c represent the read depths of LCL5 (b) and LCL6 (c), respectively (orange for HiFi and green for ONT). Tracks d to f show the GC content, repeat density, and gene density, respectively.

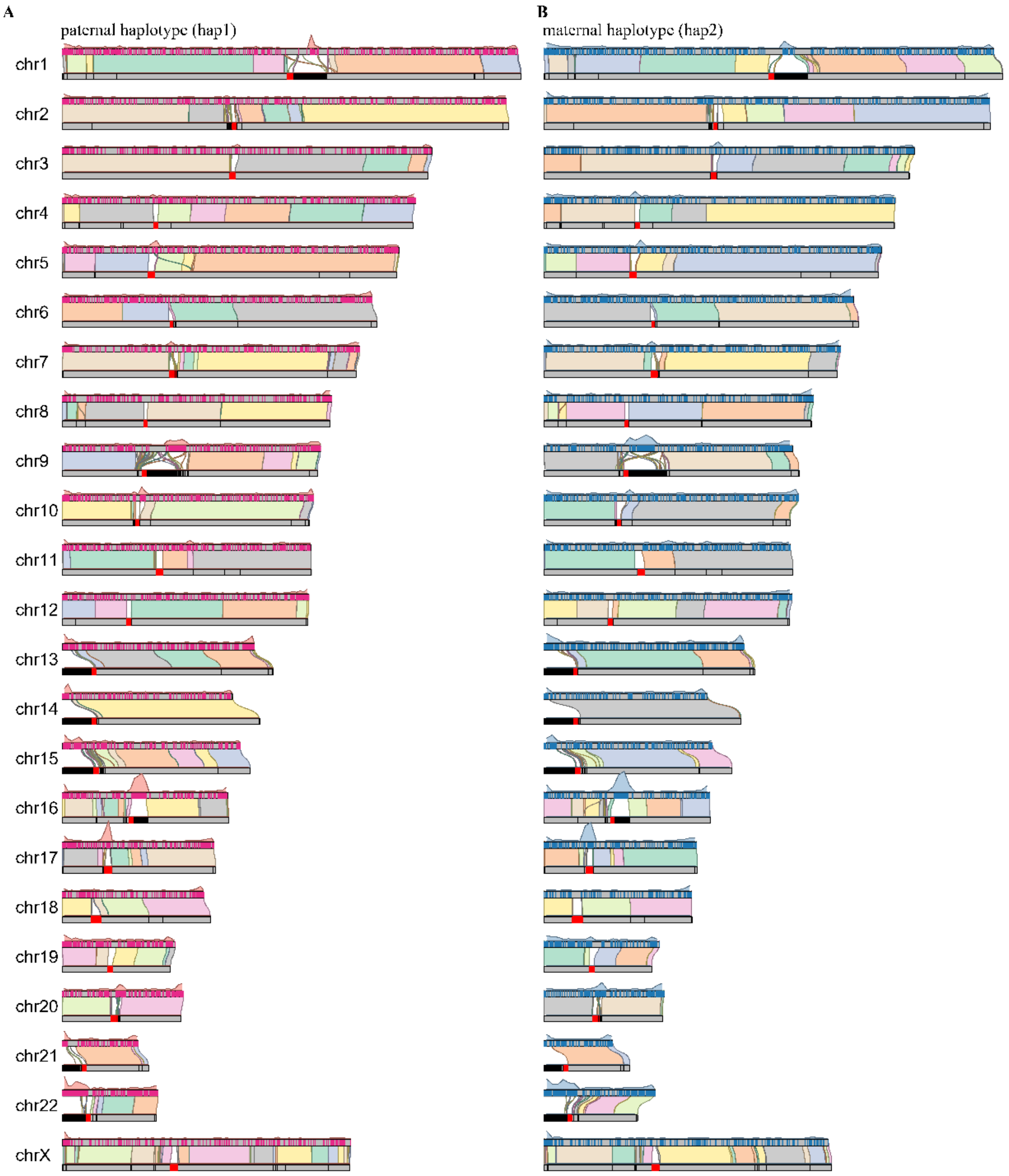

**Figure S7** Distribution of novel sequence distribution in CQ-P (left) and CQ-M (right). Alignments between haplotype (top) and GRCh38 (bottom) are represented by the links. The novel sequences of the haplotype are labeled by rectangles on the ideogram. Density plot shows the distribution of the novel sequence. Novel sequences are enriched in the centromeric and acrocentric regions of the genome.

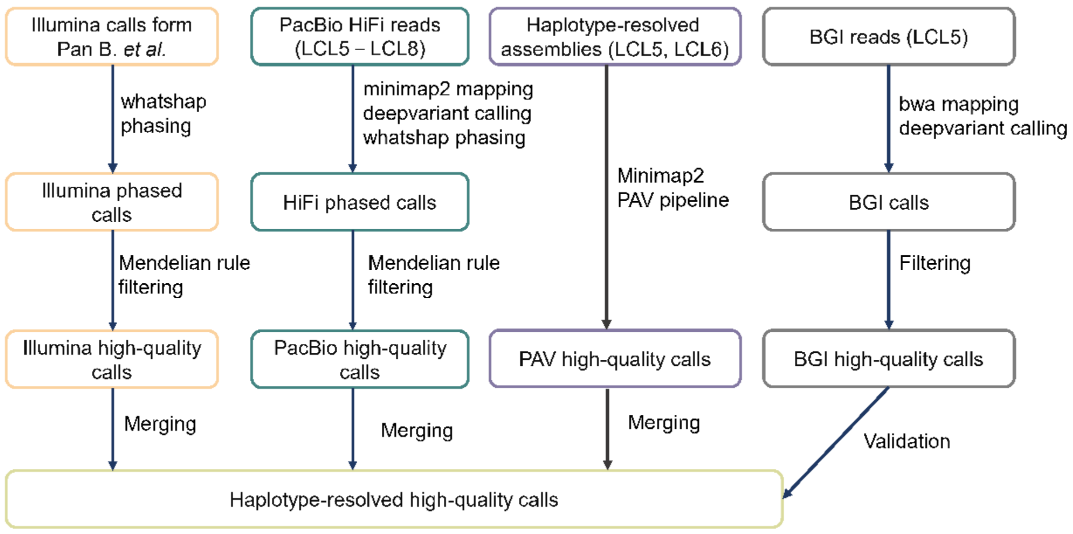

**Figure S8** SNV and Indel detection and validation pipelines.

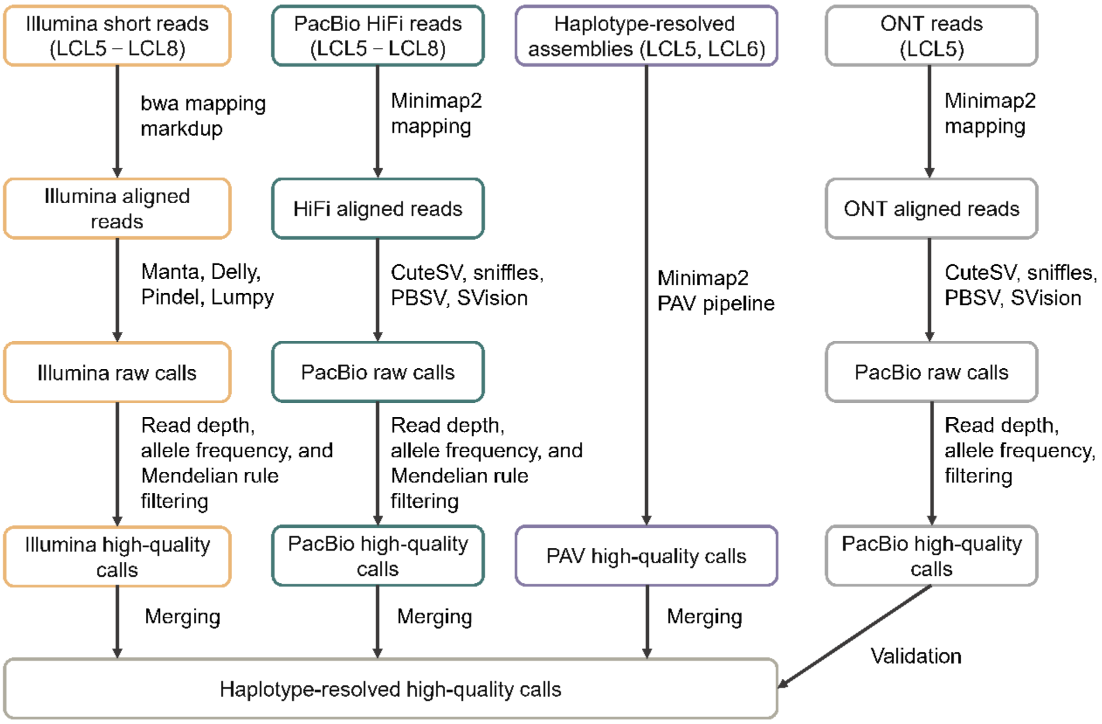

**Figure S9** Structural variant detection and validation pipelines

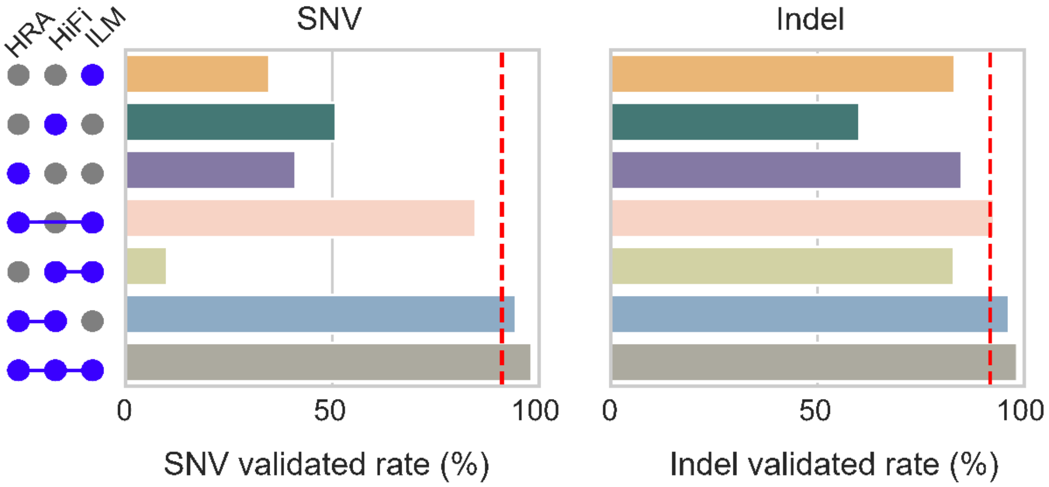

**Figure S10** Validated ratio of SNVs and Indels across different combinations of three technologies.

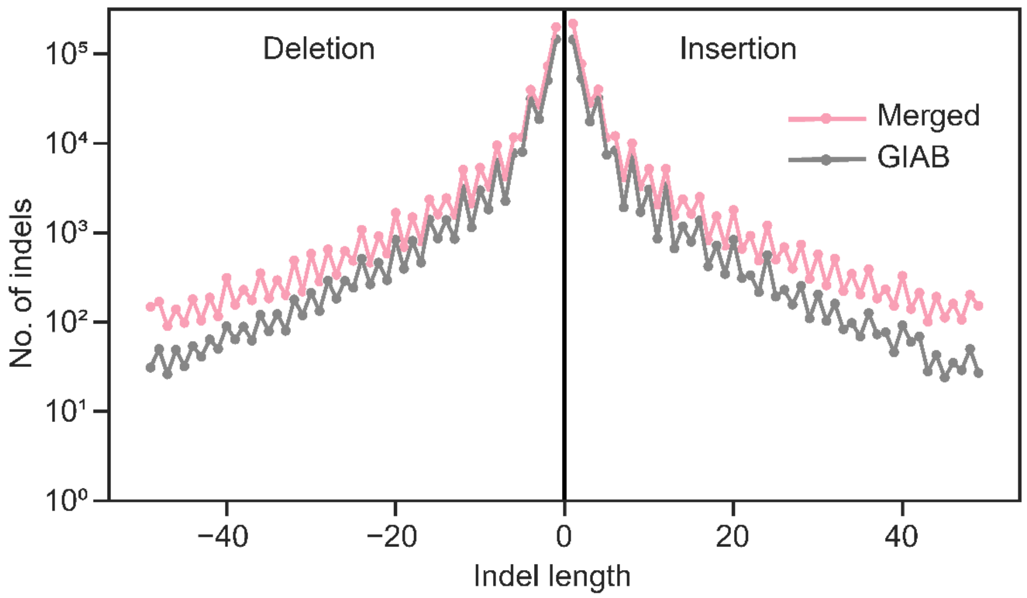

**Figure S11** Indel length distributions of HG002 and Chinese Quartet twin daughters

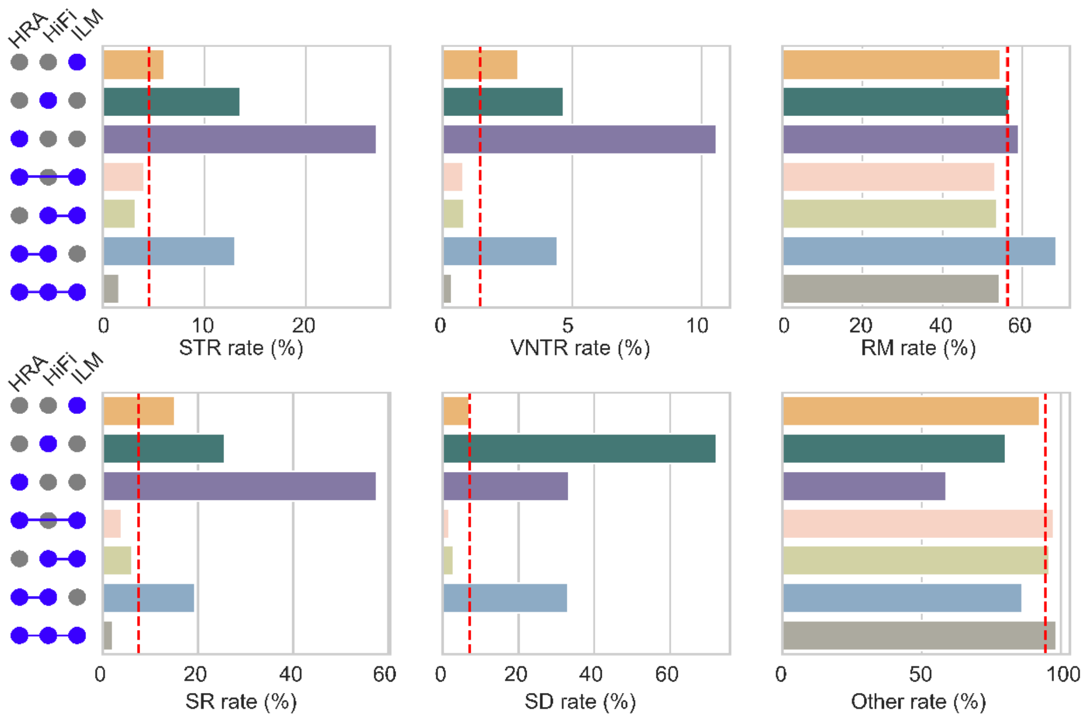

**Figure S12** SNV rate in repeat regions across different combinations of three technologies. SD, segmental duplication; SR, simple repeat; VNTR, variable number tandem repeat; STR, short tandem repeat; RM, repeat mask regions; Other, regions excluding SD, SR, VNTR, STR, and RM.

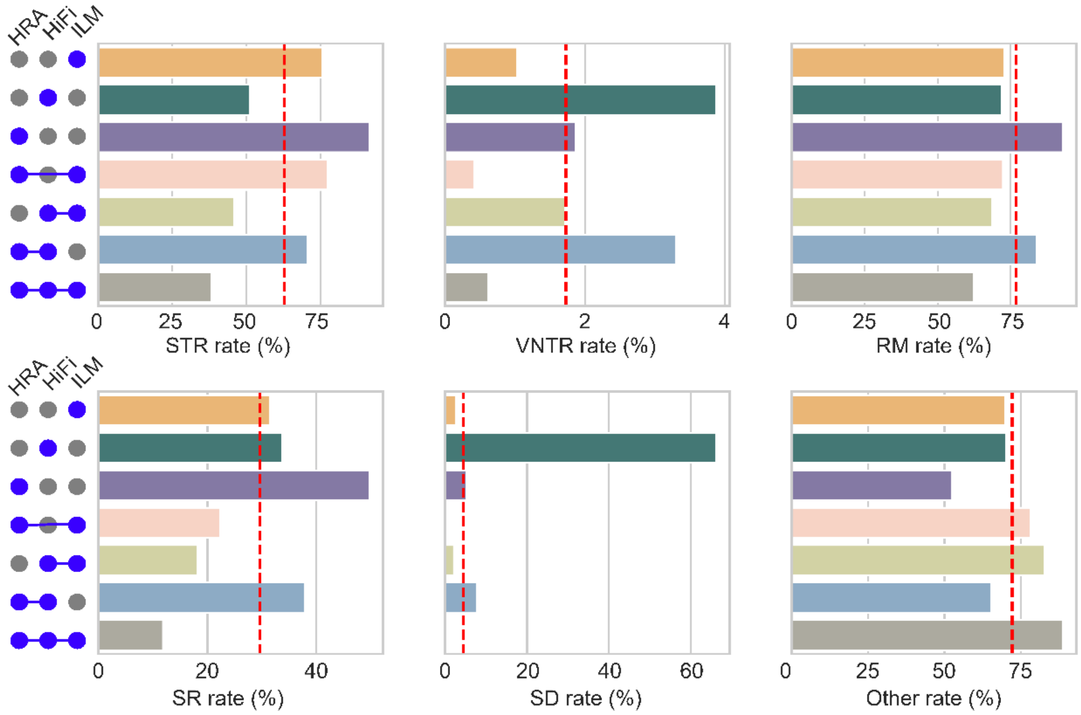

**Figure S13** Indel rate in repeat regions across different combinations of three technologies. SD, segmental duplication; SR, simple repeat; VNTR, variable number tandem repeat; STR, short tandem repeat; RM, repeat mask regions; Other, regions excluding SD, SR, VNTR, STR, and RM.

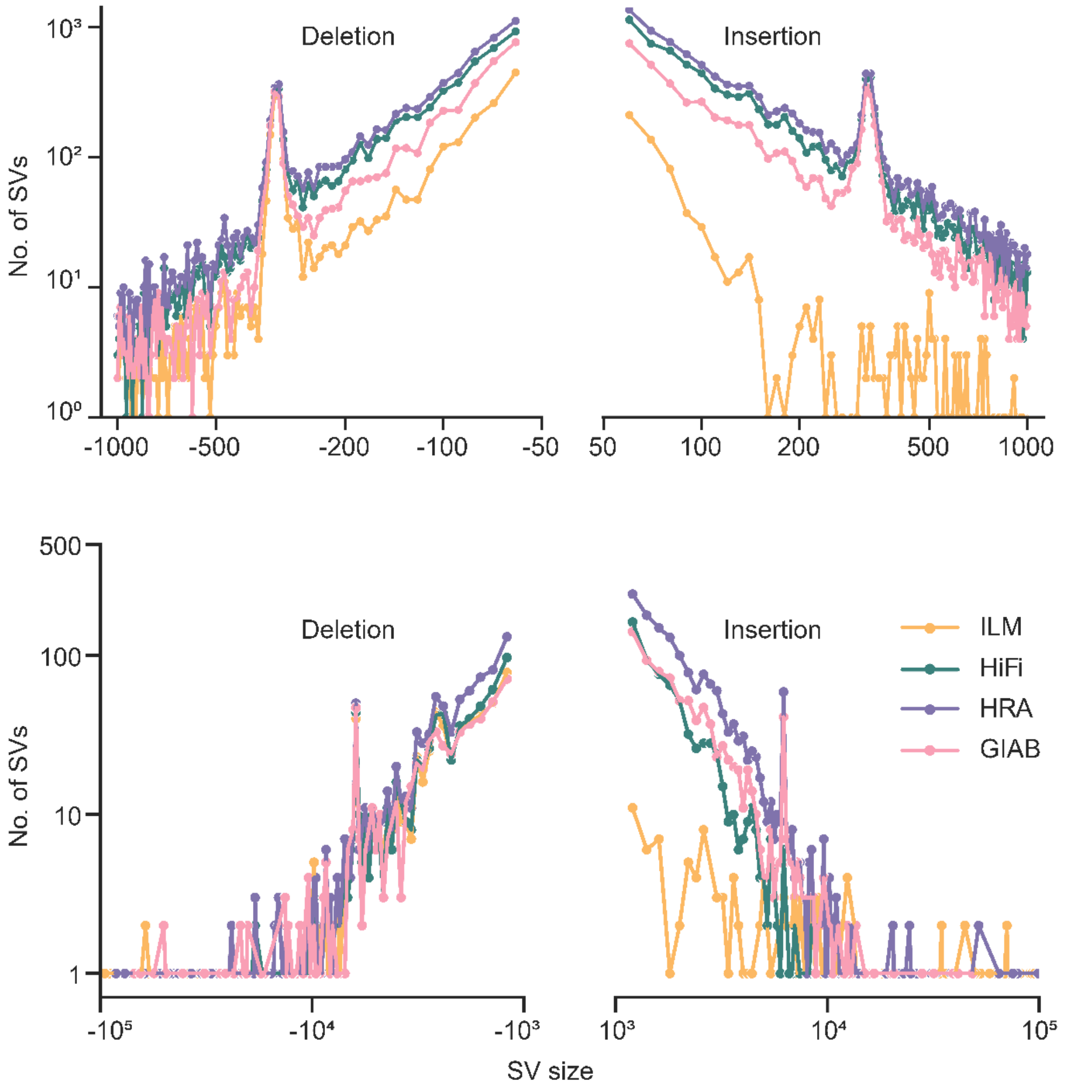

**Figure S14** SV length distributions of HG002 and three callsets of Chinese Quartet twin daughters.

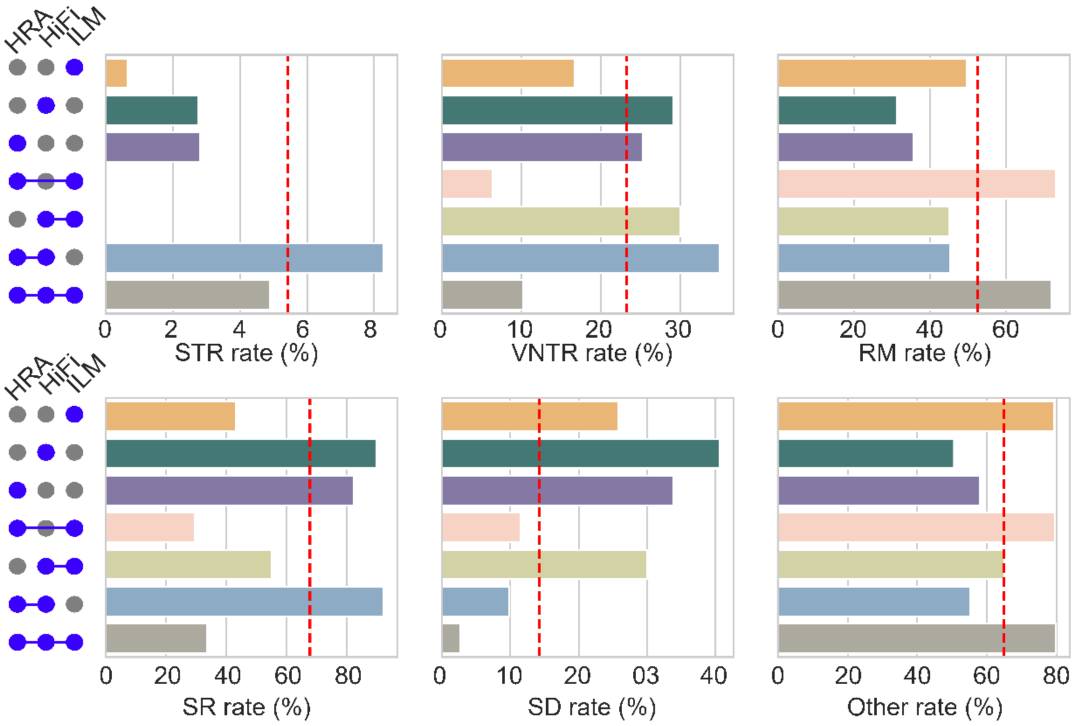

**Figure S15** Large deletion rate in repeat regions across different combinations of three technologies. SD, segmental duplication; SR, simple repeat; VNTR, variable number tandem repeat; STR, short tandem repeat; RM, repeat mask regions; Other, regions excluding SD, SR, VNTR, STR, and RM.

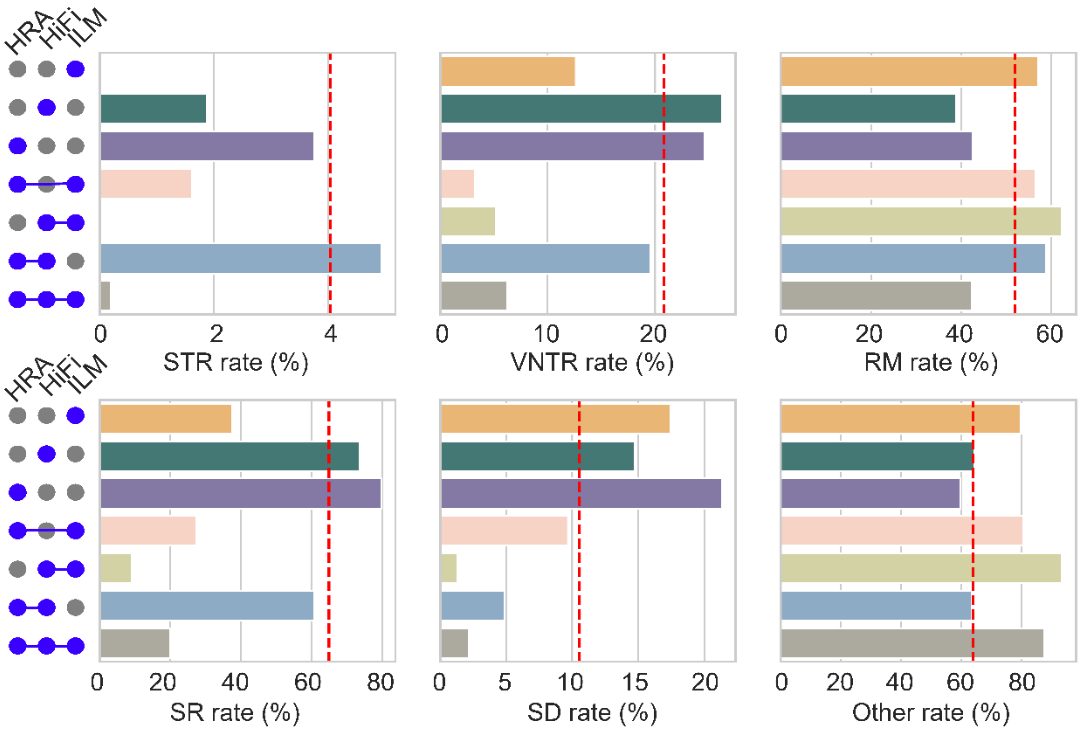

**Figure S16** Large insertion rate in repeat regions across different combinations of three technologies. SD, segmental duplication; SR, simple repeat; VNTR, variable number tandem repeat; STR, short tandem repeat; RM, repeat mask regions; Other, regions excluding SD, SR, VNTR, STR, and RM.

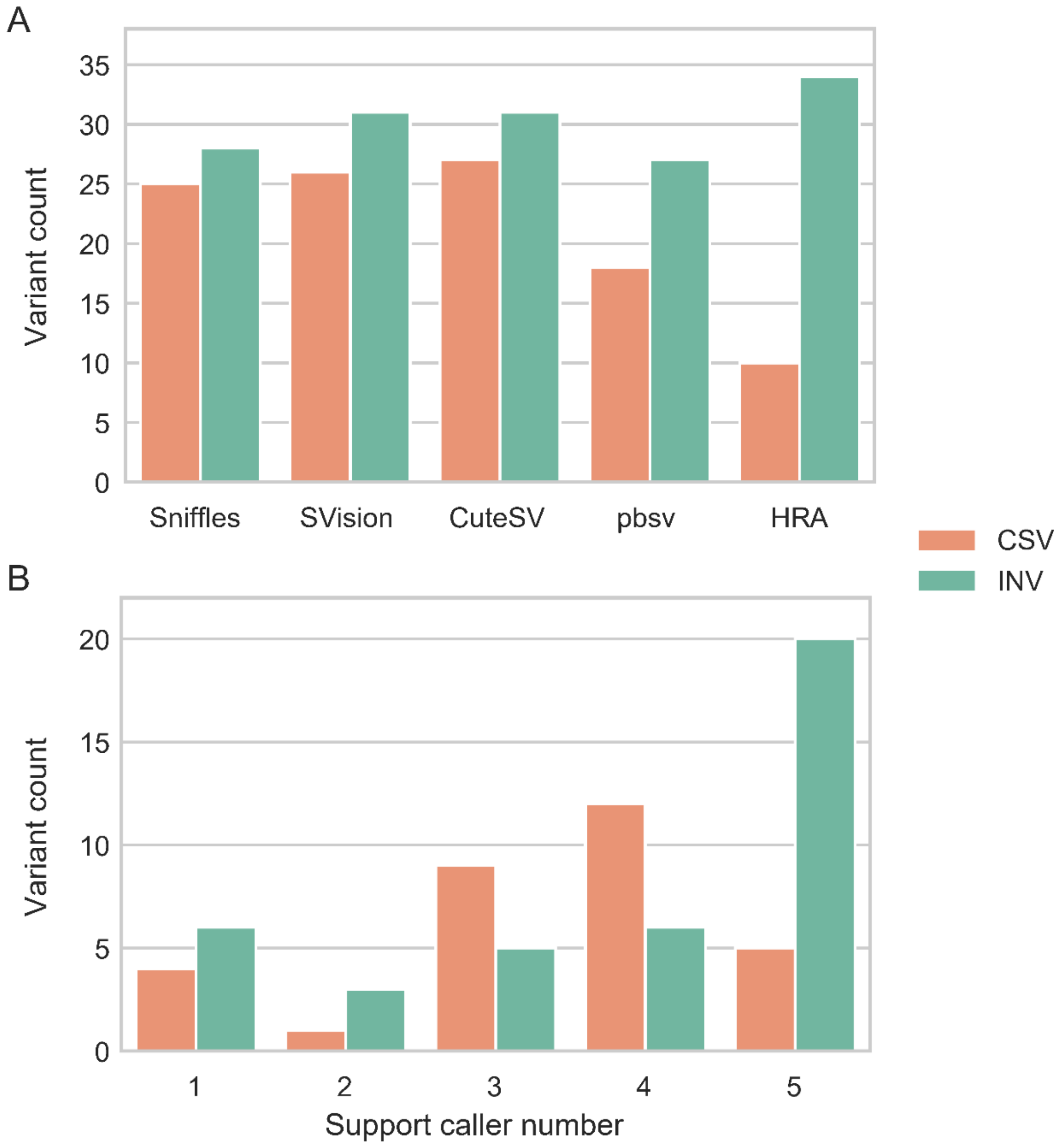

**Figure S17** Complex SVs and inversions in benchmarks. **A** Bar plot shows the number of variants discovered by different callers. **B** Bar plot shows the variant numbers of different supported callers.

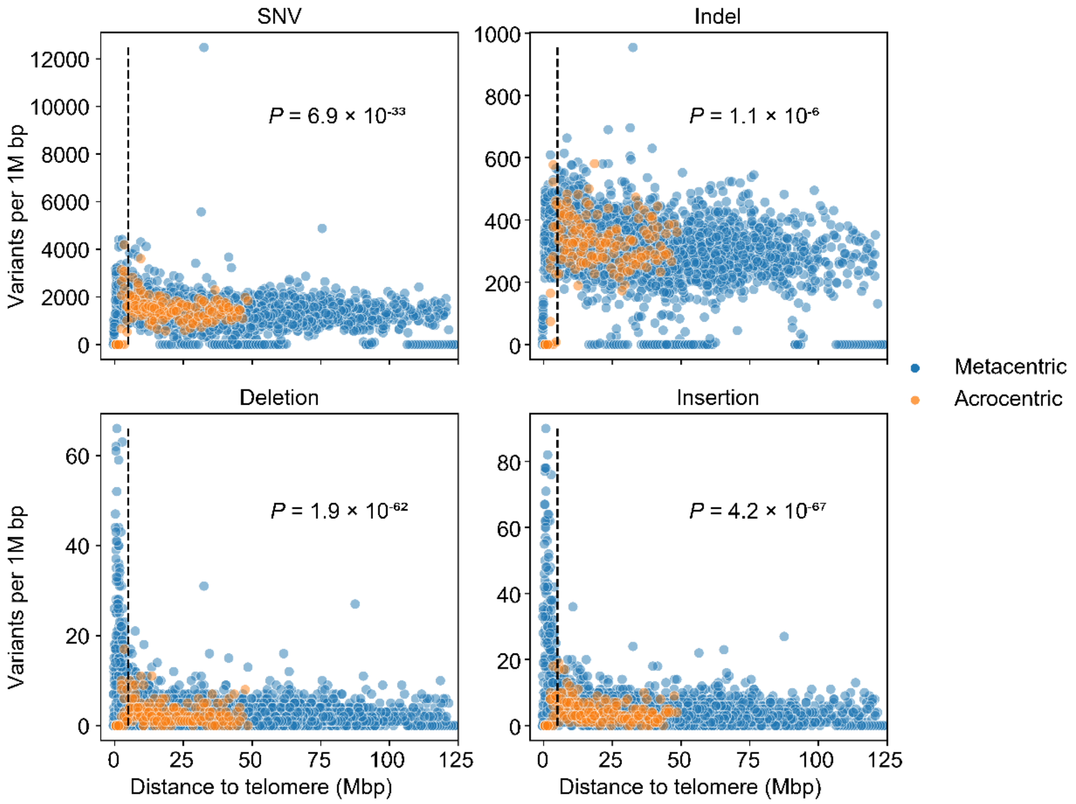

**Figure S18** Variant distribution of Chinese Quartet at the telomere. For each variant, the distance to the closest telomere of the chromosome is computed and divided into 1 Mbp bins. Variants are significantly enriched (Wilcoxon rank-sum test) within 5 Mbp of the telomere (dashed line left).

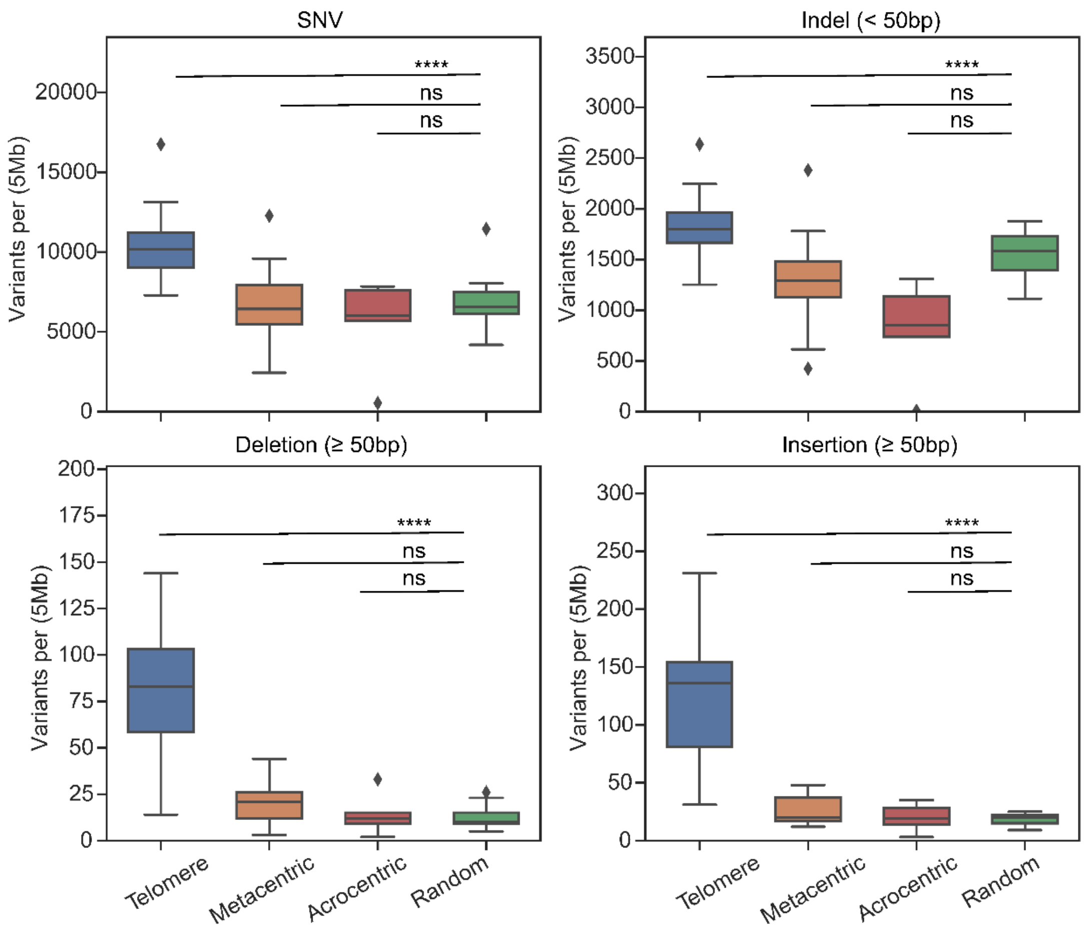

**Figure S19** Variant distribution of Chinese Quartet at telomere and centromer. The y-axis represents the number of variants located within 5 Mbp of the metacentric centromere, acrocentric centromere, and telomere. The number of variants located at telomeres is significant (Wilcoxon rank-sum test) more than other random background regions. ns, not significant; *, $P < 0.05$; **, $P < 0.01$; ***, $P < 0.001$; ****, $P < 0.0001$.

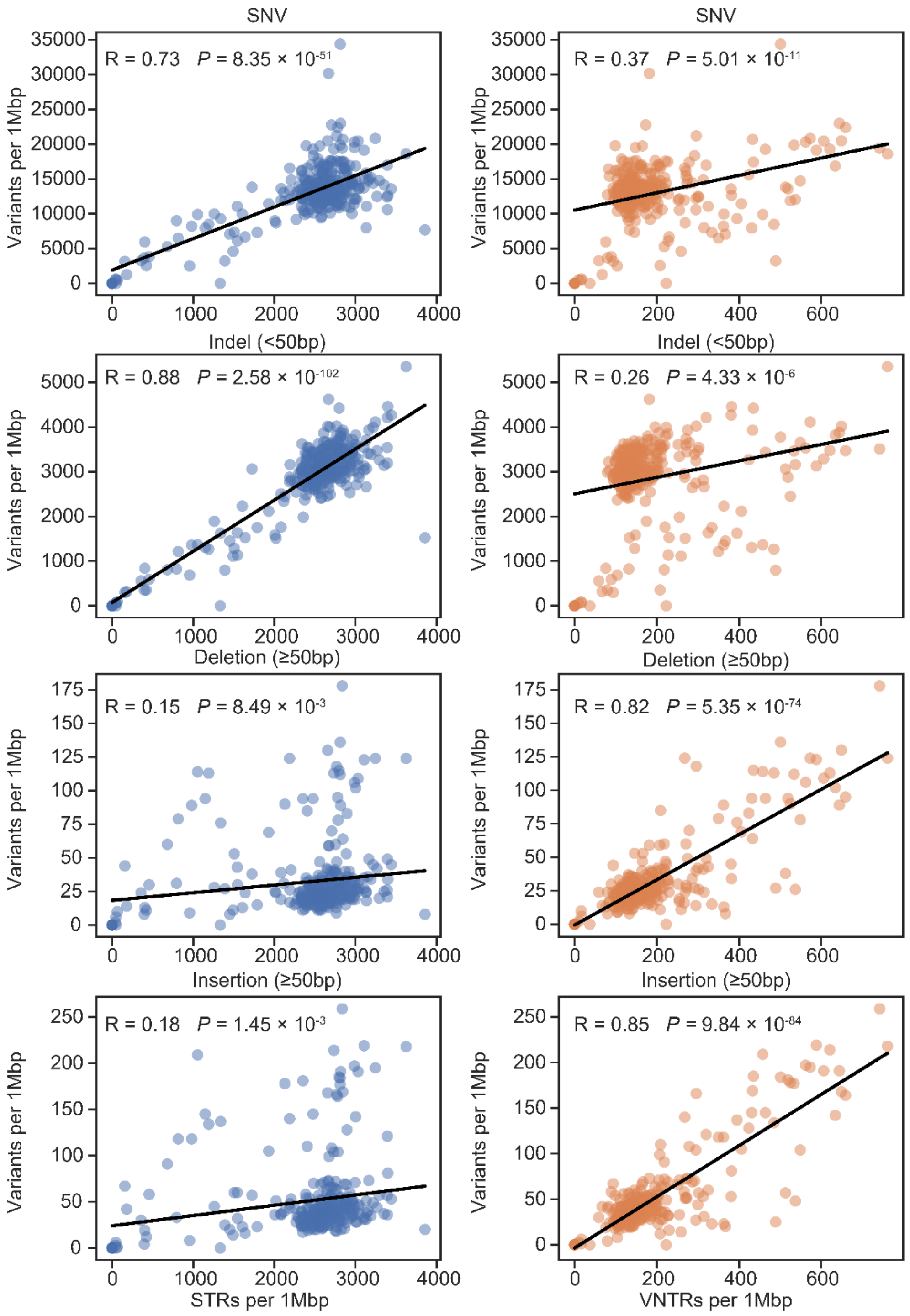

**Figure S20** STR/VNTR distribution and variant breakpoint correlation. The abundance of STRs is positively correlated with the distribution of SsNVs (R = 0.73) and Indels (R = 0.88), while VNTRs is positively correlated with structural variant (deletions, R= 0.82; insertion, R=0.85).

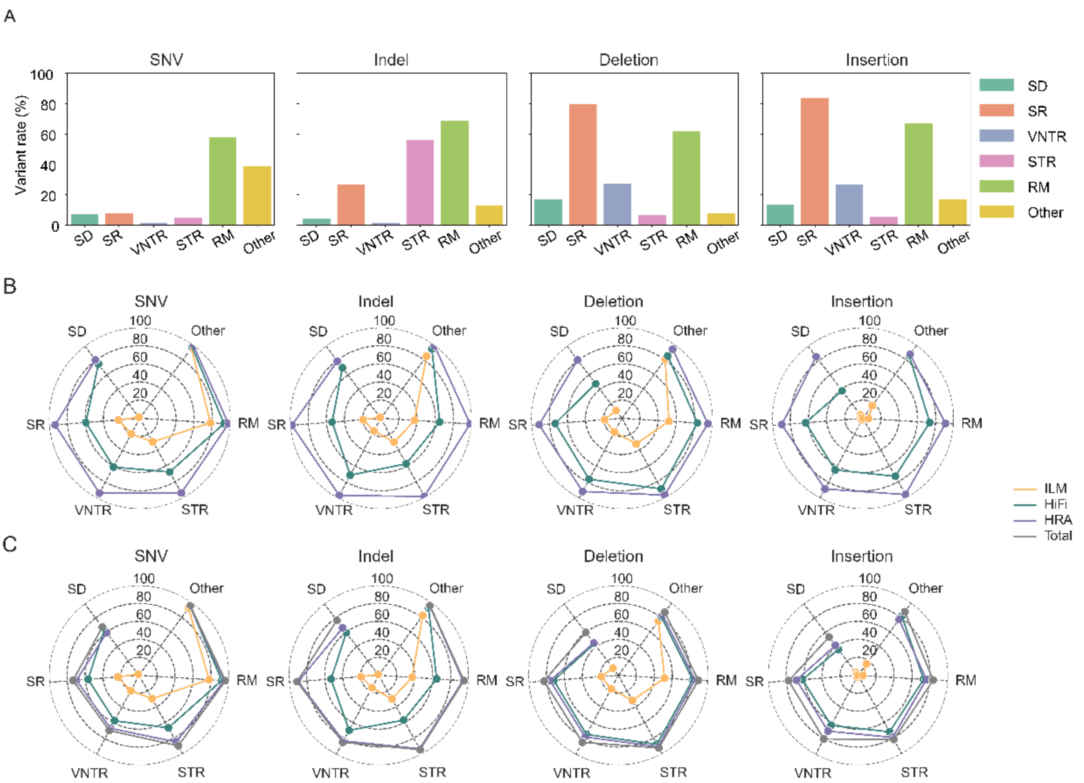

**Figure S21** Variant features in different types of regions. **A** Ratio of variants in different types of regions. **B** Radar plots show the percentage of SNVs, Indels, large deletions, and insertions detected by ILM, HiFi, and HRA in distinct regions of the genome. **C** Radar plots show the percentage of validated SNVs, Indels, large deletions, and insertions in distinct regions of the genome. SD, segmental duplication; SR, simple repeat; VNTR, variable number tandem repeat; STR, short tandem repeat; RM, repeat mask regions; Other, regions excluding SD, SR, VNTR, STR, and RM.

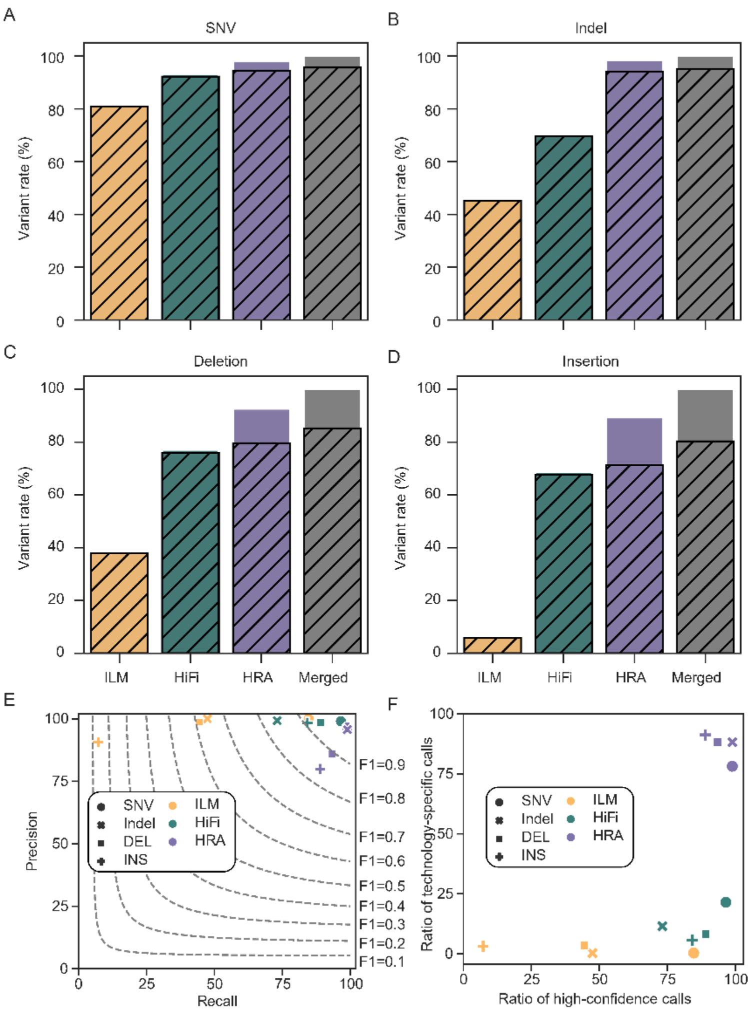

**Figure S22** Benchmark evaluation of Chinese Quartet twin daughters. **A – D** Bar plot depicts the percentage of ILM, HiFi, and HRA calls in the final benchmarks, with gray stripes representing the validated percentages by BGI or ONT reads. **E** Precision and recall of ILM, HiFi, and HRA of Chinese Quartet benchmarks. Recall of each technology is represented by the ratio of high-confidence calls this technology detects to all validated calls. Precision of each technology is defined by the percentage of validated calls in all detected calls in this technology. The dashed line denotes the F1 score of the technology. **F** Scatter plot shows the percentage of high-confidence (x-axis) and technology-specific (y-axis) calls across ILM, HiFi, and HRA.

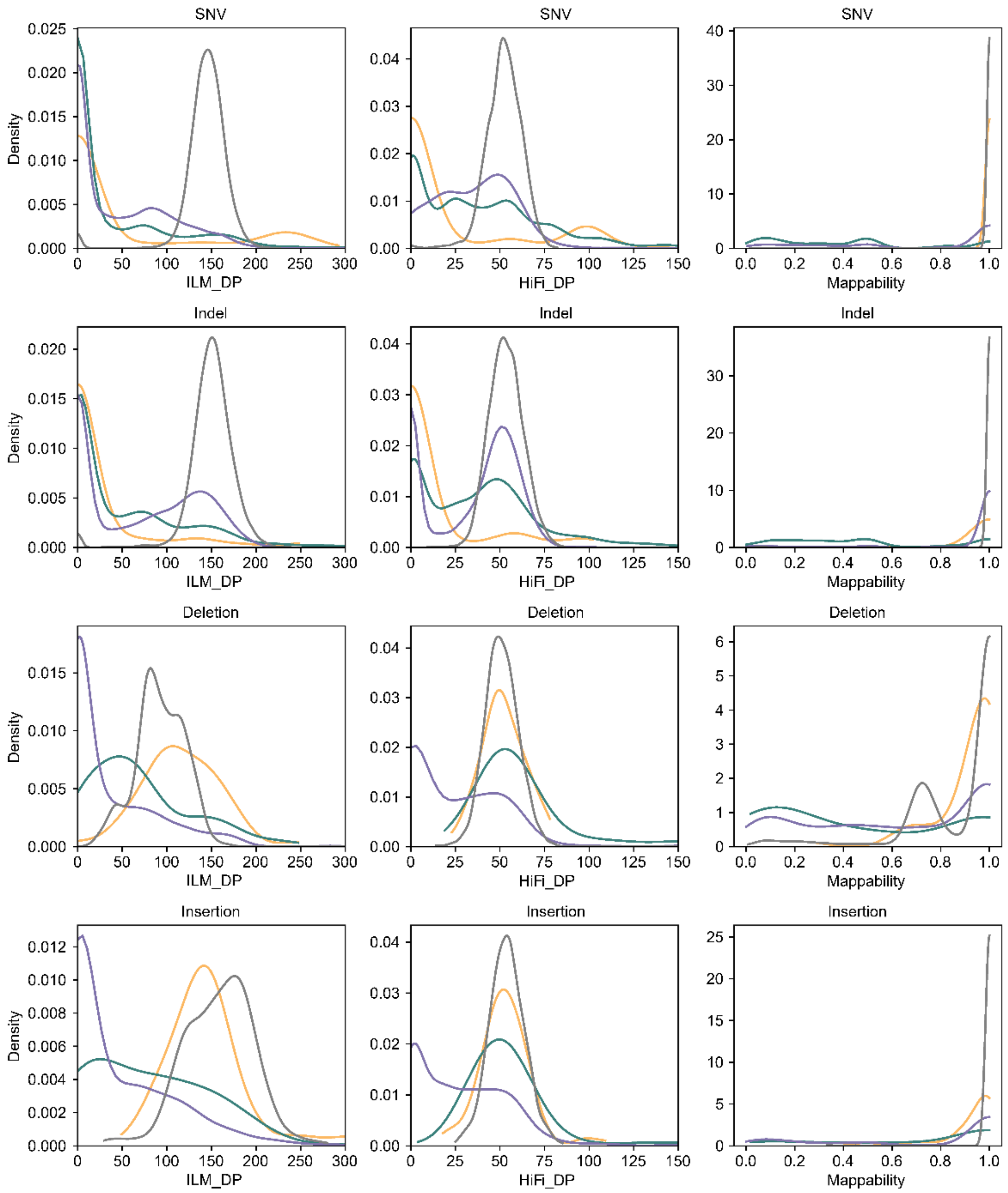

**Figure S23** The density plots show the difference in variant characteristics between high-confidence and technology-specific calls. Only reads with a mapping quality of at least 20 are used to compute read depths.
